# Dynamic adhesion device of phage OE33PA drives Gram-positive host recognition

**DOI:** 10.64898/2026.05.20.726473

**Authors:** Laura Schmitt, Amel Chaïb, Denis Ptchelkine, Eaazhisai Kandiah, Claire Le Marrec, Christian Cambillau, Adeline Goulet

## Abstract

Bacteriophages (phages) infecting Gram-positive bacteria must bind to host receptors across thick cell walls to initiate infection, yet the underlying structural mechanisms remain unclear. Here, we report cryo-electron microscopy structures of the *Oenococcus oeni* siphophage OE33PA, providing the first high-resolution view of a phage infecting this wine-industry bacterium. While the overall virion architecture is conserved, the adhesion device displays distinctive features. Its receptor-binding proteins adopt multiple orientations, revealing an intrinsically dynamic assembly. *In situ* cryo-electron tomography captures distinct conformations upon host attachment, providing rare structural insight into interactions with Gram-positive hosts. Functional assays further show that a mobile carbohydrate-binding module in the distal tail protein mediates host cell binding. Additionnally, the tape measure protein adopts a hexameric organization, updating the prevailing trimeric model in siphophages. Together, these findings reveal a dynamic adhesion device in a phage infecting Gram-positive bacteria and highlight the structural and functional diversity of phages.

## Introduction

Bacteriophages (phages), viruses that infect bacteria, are the most abundant biological entities of the biosphere. Among them, tailed double-stranded DNA phages of the class *Caudoviricetes* represent the most frequently isolated and structurally characterized group ^1^. Phages with long, non-contractile tails, commonly referred to as siphophages, constitute the largest group of cultured tailed phages ^1^.

Siphophages initiate infection through a multiprotein adhesion device at the distal end of the tail, which mediates specific host recognition and attachment ^2–4^. Following binding, genome delivery requires penetration of the bacterial cell envelope and translocation of DNA into the cytoplasm ^2^. Structural studies have shown that siphophage adhesion devices are modular assemblies typically organized around a conserved core formed by the distal tail protein (Dit) and the tail-associated lysin (Tal), with receptor-binding proteins (RBP) and accessory proteins variably attached depending on the phage ^5–14^.

Despite the availability of various high-resolution siphophage structures ^5,11–14^, the manner with which their adhesion devices interact with bacterial surfaces at the molecular and structural levels remains poorly understood. Exploring the structural diversity of phage-bacterium interactions is therefore essential for advancing our understanding of phage biology and for guiding strategies to either promote beneficial phage infections, such as for therapeutic applications, or prevent detrimental infections, such as in the food industry where Gram-positive lactic acid bacteria (LAB) are used to produce fermented foods and beverages.

This knowledge gap is especially relevant for *Oenococcus oeni*, the primary LAB responsible for malolactic fermentation in winemaking. While several structural studies have been performed on phages infecting the dairy LAB *Lactococcus lactis* and *Streptococcus thermophilus* ^7–10^, little is known about phages infecting plant-associated LAB. Lytic *O. oeni*-infecting siphophages, including the virulent Vinitor 162 and Krappator X27 as well as the ex-temperate OE33PA, that have been identified in grape must and fermentation samples can contribute to malolactic fermentation failure ^15–17^. A structural and functional understanding of the interactions between *O. oeni* and its phages is therefore essential both to advance fundamental knowledge of phage-host recognition and to better integrate phages in the ecology of this particular ecosystem and, potentially, inform strategies to prevent fermentation failures.

Here, we focused on OE33PA, whose adhesion device is predicted by AlphaFold2 to display distinct features compared with those of dairy LAB phages ^18^. To determine the structure of this device within the intact virion and investigate its role in host recognition, we used cryo-electron microscopy (cryoEM) and single particle analysis and resolved the virion architecture at high resolution, including the capsid, connector, tail, and adhesion device. While the capsid, connector, and tail resemble those of HK97-like phages, the adhesion device displays distinctive features. Notably, we observe a transition of the α-helical tape measure protein (TMP) from hexameric to trimeric symmetry within the Dit-Tal core, alongside intrinsically mobile RBP adopting multiple conformations. In addition, functional assays demonstrate that a carbohydrate-binding module (CBM) embedded within a long and highly flexible non-canonical insertion of the Dit protein mediates host cell binding. Furthermore, *in situ* cryo-electron tomography (cryoET) captures distinct conformational states of the adhesion device upon cell attachment, one of the very few structural views of cell-bound phages infecting Gram-positive hosts. Together, these findings reveal a dynamic adhesion device in a phage infecting Gram-positive bacteria and highlight the structural and functional diversity of phages.

## Results

### OE33PA cryoEM structure

OE33PA is the first phage infecting *O. oeni* to have been characterized at the genomic and physiological levels ^15^. Its structural genes form a conserved module organized according to the canonical head-tail arrangement of LAB siphophages (Fig. 1a) ^6–8,19,20^. OE33PA exhibits the typical siphophage architecture, consisting of an icosahedral capsid connected through a head-tail connector to a long, flexible, non-contractile tail that terminates in a distal adhesion device (Fig. 1b). We determined the structures of the major virion components, including the capsid, connector, tail tube and adhesion device, representing the complete ∼330 nm-long phage particle (Fig. 1c; Supplementary Fig. 1, 2 and Table 1). The majority of phage particles contained densely packaged genomic DNA (Fig. 1b), indicating that the structure corresponds to the mature, pre-adsorption state of the virion.

**Fig. 1.**
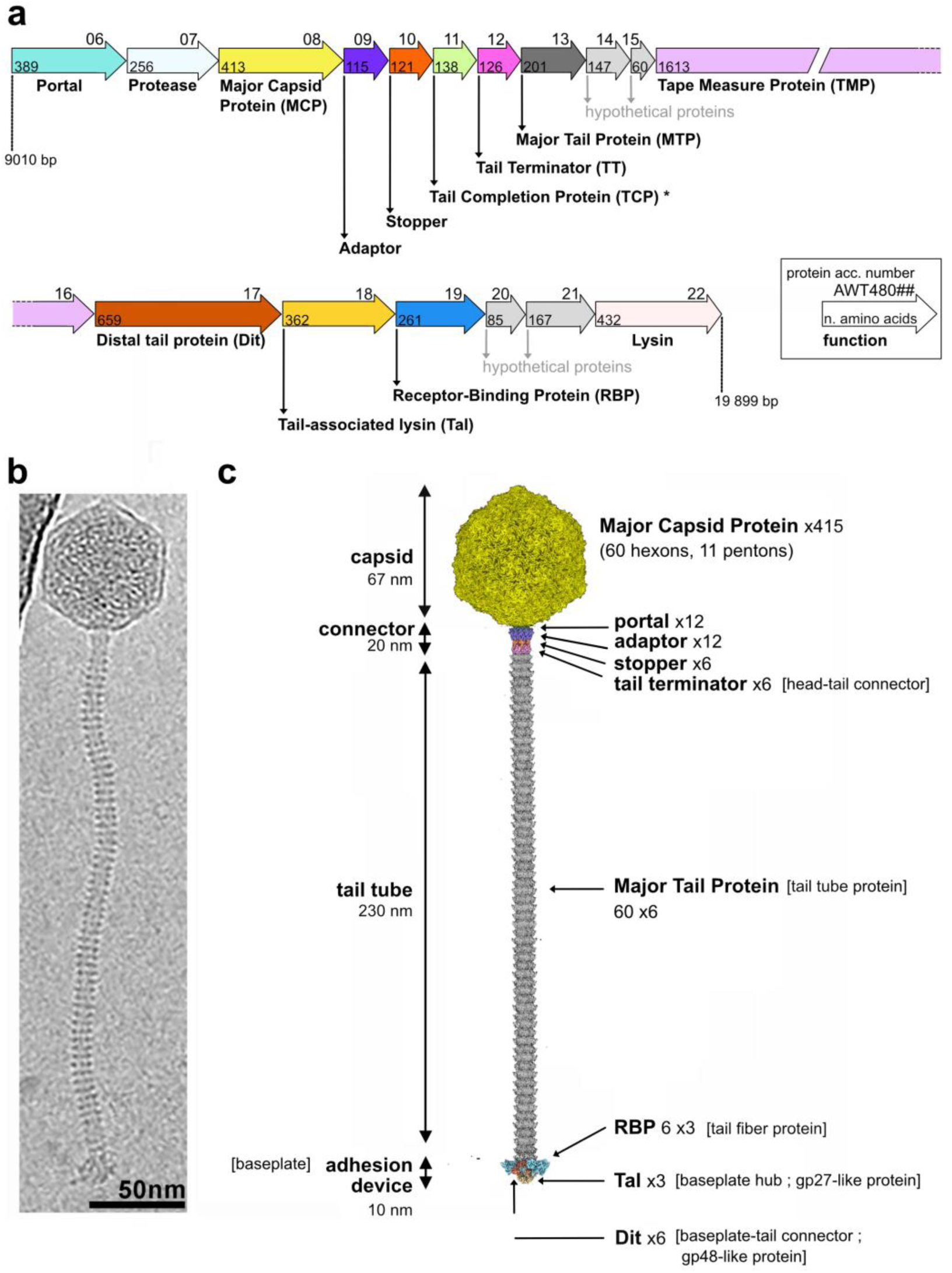
OE33PA cryoEM structure. **a**. Schematic representation of the genomic region spanning from the *portal* to the *lysin* genes. The inset explains the information provided above, within and below each arrow, corresponding to the protein accession number (only the last two digits are indicated), the number of amino acids, and the functional annotation, respectively. The asterisk indicates that the protein AWT48011 is a candidate Tail Completion Protein based on its AlphaFold3-predicted structure (Supplementary Fig. 9). **b.** CryoEM micrograph of an individual virion. **c.** 3D reconstructions of the OE33PA capsid, connector, tail and adhesion device assembled into a complete virion (color code as in panel a). The tail was assembled from the 3D reconstruction of two MTP rings to match the 60 MTP rings counted in the micrograph shown in b. Common alternatives names for proteins are provided in brackets.

### Capsid

The OE33PA capsid was reconstructed with imposed icosahedral symmetry to an overall resolution of 2.8 Å (Fig. 2a; Supplementary Fig. 1, 4, 6 and Table 1). The ∼67 nm-diameter particle adopts a T = 7 *laevo* icosahedral geometry and encloses concentric layers of dsDNA (Fig. 2b). The capsid lattice is composed of 60 hexons and 11 pentons formed by the Major Capsid Protein (MCP), for a total of 415 MCP subunits. No additional cement or decoration proteins were observed.

**Fig. 2.**
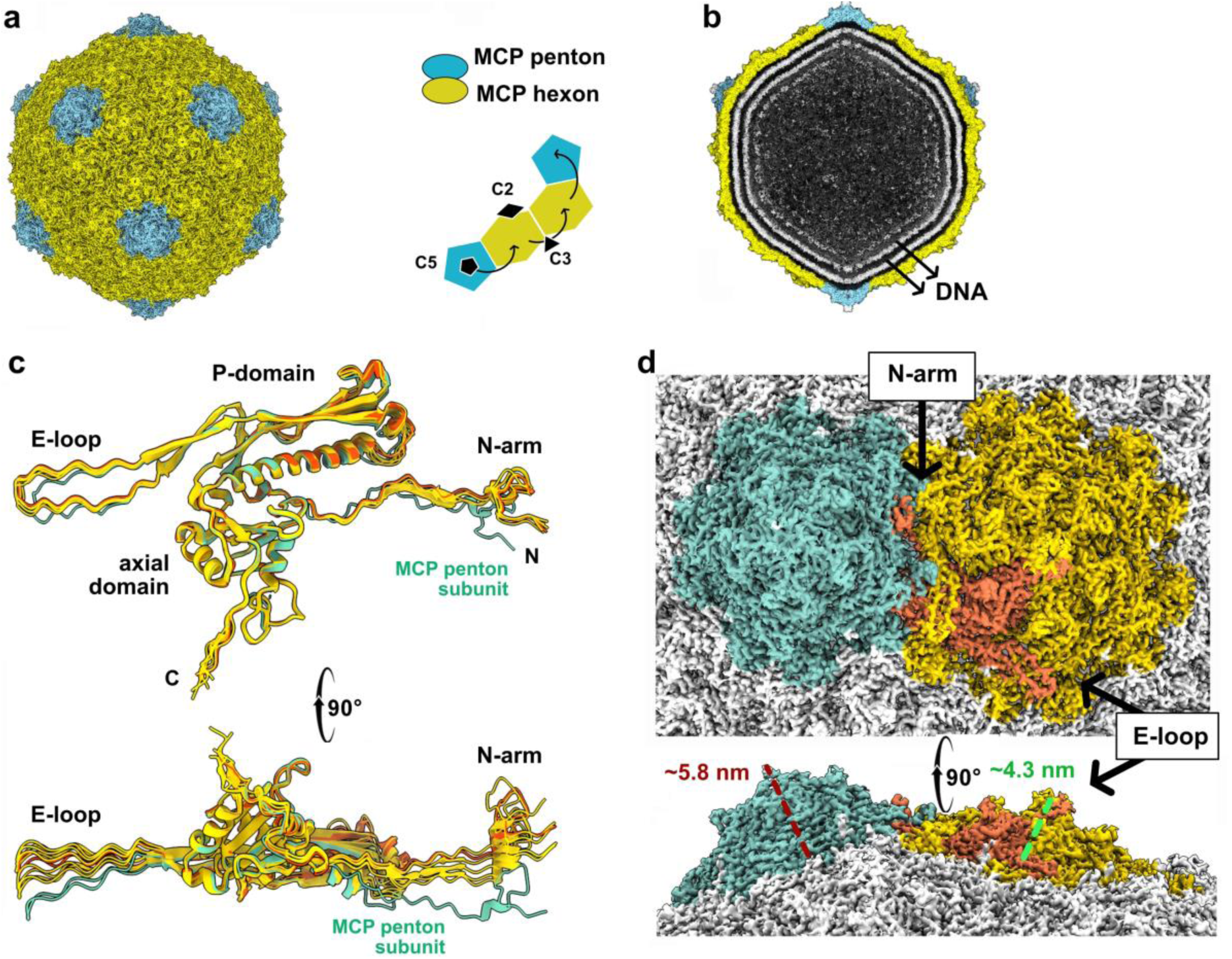
Capsid structure. **a**. 3D reconstruction of the capsid with imposed icosahedral symmetry. The MCP hexons and pentons are shown in yellow and blue, respectively. The schematic highlights the T = 7 *laevo* icosahedral geometry. **b.** Cut-away view of the capsid reconstruction in the same orientation as in a, revealing concentric rings of the phage dsDNA genome. **c**. Othogonal views of ribbon representations of the superimposed MCP subunits from a capsid asymmetric unit, including 6 subunits from the hexon (yellow and orange), and one subunit from the penton (blue). **d.** Othogonal views of 3D reconstruction focused on one penton (blue) and one hexon (yellow, orange). One hexon MCP subunit is orange as in c.

The OE33PA MCP adopts the canonical HK97 fold ^21^, consisting of an N-terminal arm (N-arm, residues 116-149), an extended loop (E-loop, residues 163-201), a peripheral domain (P-domain, residues 203-221, 335-381), and an axial domain (residues 249-348, 389-404) (Fig. 2c). Structural similarity searches using Foldseek ^22^ identified significant matches to the MCP of the *Rhodobacter capsulatus* gene transfer agent (GTA) ^23^ and phage HK97 ^21^ (Supplementary Table 2 and Fig. 7), supporting its classification within the HK97-like lineage. No density was observed before residue M116, indicating proteolytic removal of a maturation domain. This processing is consistent with the maturation pathway described for HK97-like phages ^24^. The N-arm and E-loop extend in opposite directions from each MCP subunit, forming extensive interactions with the two adjacent MCP subunits within the same capsomer (∼ 2400 Å^2^ buried surface area) and bridging to MCP subunits of neighboring capsomers (∼ 340 Å^2^ buried surface area for the N-arm) (Fig. 2d). These long-range, intertwined interactions form an interconnected network that stabilizes the capsid shell ^25^.

The shell thickness varies from ∼2.5 nm at its thinnest regions to ∼4.3 nm and ∼5.8 nm at the centers of hexons and pentons, respectively, where the MCP C-termini form protruding knobs at the capsid surface (Fig. 2a, d). Superposition of MCP subunits within the asymmetric unit, comprising one penton subunit and one hexon, reveals pronounced conformational differences. In the penton subunit, the N-arm and E-loop are rotated by approximately 20° and 30°, respectively, relative to their hexon counterparts (Fig. 2c). This arch-shaped conformation accounts for the increased protrusion of pentons compared to hexons and reflects the conformational plasticity required for the MCP to occupy pentameric versus hexameric environments within the T = 7 lattice.

### Connector, connector-capsid interface and connector-DNA assembly

The connector replaces one MCP penton at a single fivefold vertex of the capsid and links the capsid to the tail. It is composed of a dodecameric portal protein embedded within the capsid, followed sequentially by a dodecameric adaptor, a hexameric stopper, and a hexameric tail terminator (TT), the latter engaging the tail tube formed by stacked ring hexamers of the major tail protein (MTP) (Fig. 1c, 3a). We reconstructed the entire connector in the context of the capsid vertex, including the five surrounding MCP hexons, without imposing symmetry, to an overall resolution of 3.6 Å (Supplementary Fig. 1, 4 and Table 1). Localized reconstructions imposing C12 symmetry for the portal-adaptor region, and C6 symmetry for the stopper-tail terminator complex bound to one MTP hexamer, were refined to 2.6 Å and 2.8 Å resolution, respectively, and used for model building (Supplementary Fig. 1, 4, 6 and Table 1). We also reconstructed the connector and the first MTP ring to an overall resolution of 3.0 Å imposing C6 symmetry to examine protein-protein interfaces within the entire complex (Fig. 3a; Supplementary Fig. 1, 4 and Table 1). The OE33PA connector forms a ∼20 nm-long densely packed assembly with external diameters of ∼15 nm at the portal and ∼7.5 nm at the stopper and TT (Fig. 3a).

**Fig. 3.**
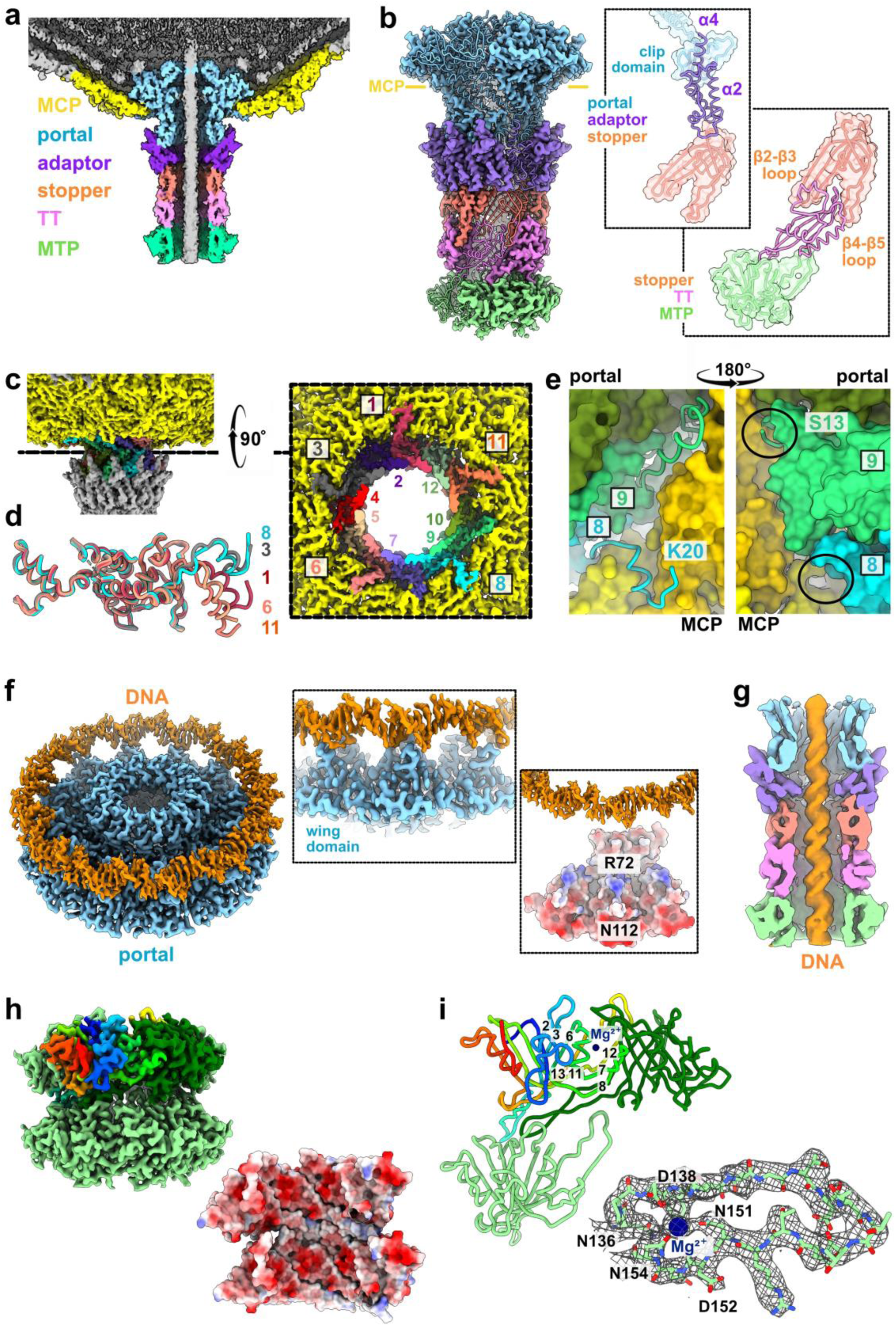
**Connector and tail structures**. **a**. Cut away view of the reconstruction of the capsid-connector assembly (C1 symmetry). **b.** Reconstruction of the connector (C6 symmetry imposed) and ribbon representation of portal (x2), adaptor (1), stopper (x1) tail terminator (x1) and MTP (x1) subunits (the corresponding densities are not shown for clarity) in the same orientation as in a. The yellow lines indicate the position of the capsid. The black boxes show surface and ribbon representations of protein-protein interfaces. **c**. Orthogonal views of the reconstruction of the capsid-portal assembly (C1 symmetry). The capsid is yellow and the 12 portal subunits are numbered and shown in different colors. **d.** Ribbon representations of the superimposed five portal subunits with N-terminal extensions. **e**. Close-up views of the interface between two MCP subunits (light and dark yellow) and two portal subunits (blue and green). The black circles highlight the absence (MCP subunit 8) and the presence (MCP subunit 9) of the N-terminal extension starting at S13. **f**. Reconstruction of the portal-adaptor assembly (C12 symmetry imposed) with DNA. The black boxes show a close-up view of the DNA-portal interface (with only three portal subunits shown for clarity.) next to a split view of the DNA and electrostatic surfaces of portal subunits. **g.** 3D class of the connector with DNA inside the central channel and extending to the first MTP ring, shown in the same orientation as in a. **h.** Reconstruction of two stacked MTP rings (C6 symmetry imposed) and electrostatic surface of the MTP rings. Some MTP subunits are not shown for clarity. **i.** Ribbon representations of three MTP subunits shown in the same orientation as in h. Some β-strands are labelled. In the close-up view of the cation-binding site in the β12-β13 loop, the density is shown as a mesh, the potential Mg^2+^ ion as a sphere, and some coordinating residues are labelled.

The portal protein adopts the canonical portal fold comprising the wing, stem clip and crown domains (Supplementary Fig. 8a). Structural similarity searches identify strong homology with the portals of the *R. capsulatus* GTA and phage HK97 (Supplementary Table 2 and Fig. 7). Approximately half of the portal dodecamer, including the wing and crown domains, is inserted within the capsid, whereas the stem and clip domains extend outward (Fig. 3a, b).

The N-termini of the twelve portal subunits, which mediate interactions with the capsid shell, adopt distinct conformations. Five portal subunits display well-resolved N-terminal motifs (residues 20-43) that extend toward the center of the five surrounding MCP hexons (Fig. 3c,d; Supplementary Fig. 8b). Each motif comprises a short linker (residues 20-23) that runs along a β-sheet of one MCP subunit, a short α-helix (residues 25-32) inserted into a shallow groove at the interface between two MCP subunits, and an extended segment (residues 33-43) connecting to the portal main structure and threading between portal and MCP proteins (Fig. 3e). The uneven distribution of these N-terminal extensions within the dodecameric portal assembly, together with their distinct orientations toward surrounding MCP subunits (Fig. 3e, f), provides the molecular basis for accommodating the C5-C12 symmetry mismatch at the capsid-connector interface. The remaining seven portal N-termini are less resolved and are modeled starting between residues P37 and I39, primarily as extended segments at the portal-capsid interface. However, one of these portal subunits displays an extended N-terminal stretch beginning at S13 that traverses the capsid shell at the interface between an MCP subunit and the portal wing domain, reaching the capsid interior, where it may further stabilize the asymmetric capsid-connector interface (Fig. 3e).

The adaptor, structurally similar to its counterparts in HK97 and the *Bacillus subtilis* phage SPP1 ^26,27^ (Supplementary Table 2 and Fig. 7), consists of four α-helices (residues 2-107). At one end, the adaptor’s C-terminal α-helix forms the primary interaction with the clip domain of the portal subunit directly above, while the C-terminal tip of α-helix 2 contacts, to a lesser extent, the clip domain of the adjacent portal subunit (Fig. 3b). At the opposite end, the adaptor’s N-terminal α-helix, together with the N-terminal tip of its second α-helix and the intervening loop, establishes contacts with the hexameric stopper ring (Fig. 3b). The C12-C6 symmetry mismatch at the adaptor-stopper interface is resolved by two adaptor subunits interacting with a single stopper subunit.

The stopper adopts a canonical U-shaped fold composed of two three-stranded β-sandwiches connected by extended β2-β3 and β4-β5 loops (Fig. 3b, Supplementary Fig. 8a), resembling the stoppers of GTA and SPP1^23,27^ (Supplementary Table 2 and Fig. 7). At one end, the β-sandwich surfaces contact the adaptor ring, whereas at the opposite end the long loops interact with the TT hexamer (Fig. 3b,f). The TT is formed by a central five-stranded antiparallel β-sheet flanked by α-helices (Fig. 3b, Supplementary Fig. 8a) and resembles the TT of SPP1 and the *Agrobacterium tumefaciens* phage Milano ^28,29^ (Supplementary Table 2 and Fig. 7). Each stopper subunit embraces one TT subunit via its extended loops, which also contact the first MTP ring, anchoring the tail to the capsid (Fig. 3b).

Directly above the portal within the capsid, a closed density consistent with dsDNA, with discernible major and minor grooves, is located near the portal wing domains (Fig. 3f, Supplementary Fig. 3, 8c). The portal residues R72 and N112 are well positioned to contact the DNA (Fig. 3f). Moreover, localized 3D classifications with a mask applied to the density within the central channel of the connector revealed a continuous DNA molecule extending into the directly adjoining MTP ring (Fig. 3g, Supplementary Fig. 1, 4, 8d,e).

### Tail tube

The tail tube, formed by 60 stacked MTP hexamers, contacts the TT ring at the tube’s proximal end and the Dit of the adhesion device at the tube’s distal end (Fig. 1b, 1c). Due to the inherent flexibility of siphophage tail tubes, achieving atomic resolution is challenging. To overcome this, we reconstructed the two MTP rings immediately above the adhesion device, which form a straight segment, using localized 3D classification and refinement with C6 symmetry imposed, yielding an overall resolution of 2.9 Å (Fig. 3h, Supplementary Table 1 and Fig. 2, 5).

Each MTP adopts the canonical fold, comprising two central β-sandwiches and two α-helices exposed on the same side, with three long loops protruding from the central core (β2-β3 loop, β3-β6 loop, β12-β13 loop) (Fig. 3h). This architecture closely resembles the MTP structures of the *Pseudomonas aeruginosa* myophage E217 ^30^ and the *E.coli* siphophage T5 ^31^ (Supplementary Table 2 and Fig. 7).

Intra-hexamer interactions involve the β7-β8 loop and the β12-β13 loop, the latter forming an inter-subunit β-sheet with the β3 strand on the adjacent MTP subunit (Fig.3h). Inter-hexamer interactions are primarily mediated by the β2-β3, β3-β6, and β12-β13 loops which likely contribute to the structural flexibility of the tail tube (Fig. 3h). The 42 Å-wide internal channel is predominantly negatively charged (Fig. 3h), which may facilitate genome ejection.

A unique feature of the OE33PA MTP is the presence of a density in the 3D reconstruction, located between the β12 and β13 strands (Fig. 3i). As the surrounding residues are predominantly negatively charged and the phage purification buffer contains Mg^2+^, we tentatively assigned this density to a Mg²⁺ ion. The ion appears to be coordinated by the main chain carbonyl oxygens of D138 and D152 and the side chain carbonyl oxygens of N136, D138 and N151. However, the resolution of our reconstructions does not allow us to reliably distinguish Mg^2+^ from other possible cations, such as Ca^2+^, based on their coordination geometry; therefore, the identity of the ion cannot be definitely established. This ion may contribute to stabilizing the long β12-β13 loop, which is itself involved in intra- and inter-hexamer interactions.

### Adhesion device and TMP

We reconstructed the complete adhesion device bound to the last MTP ring without imposing symmetry, achieving an overall resolution of 3.1 Å (Fig. 1c, 4c; Supplementary Table 1 and Fig. 2, 5). It comprises a hexameric Dit ring attached to the last MTP ring, a Tal trimer connected to the opposite side of the Dit ring, and RBP trimers positioned at the periphery of the Dit-Tal core (Fig. 4c). Notably, the TMP spans the Dit-Tal channel and partially extends into the proximal MTP ring, forming a hexameric bundle within the Dit ring. However, within the Tal cavity, this arrangement transitions to a three-helix bundle formed by alternating TMP subunits (Fig. 4d,k).

**Fig. 4.**
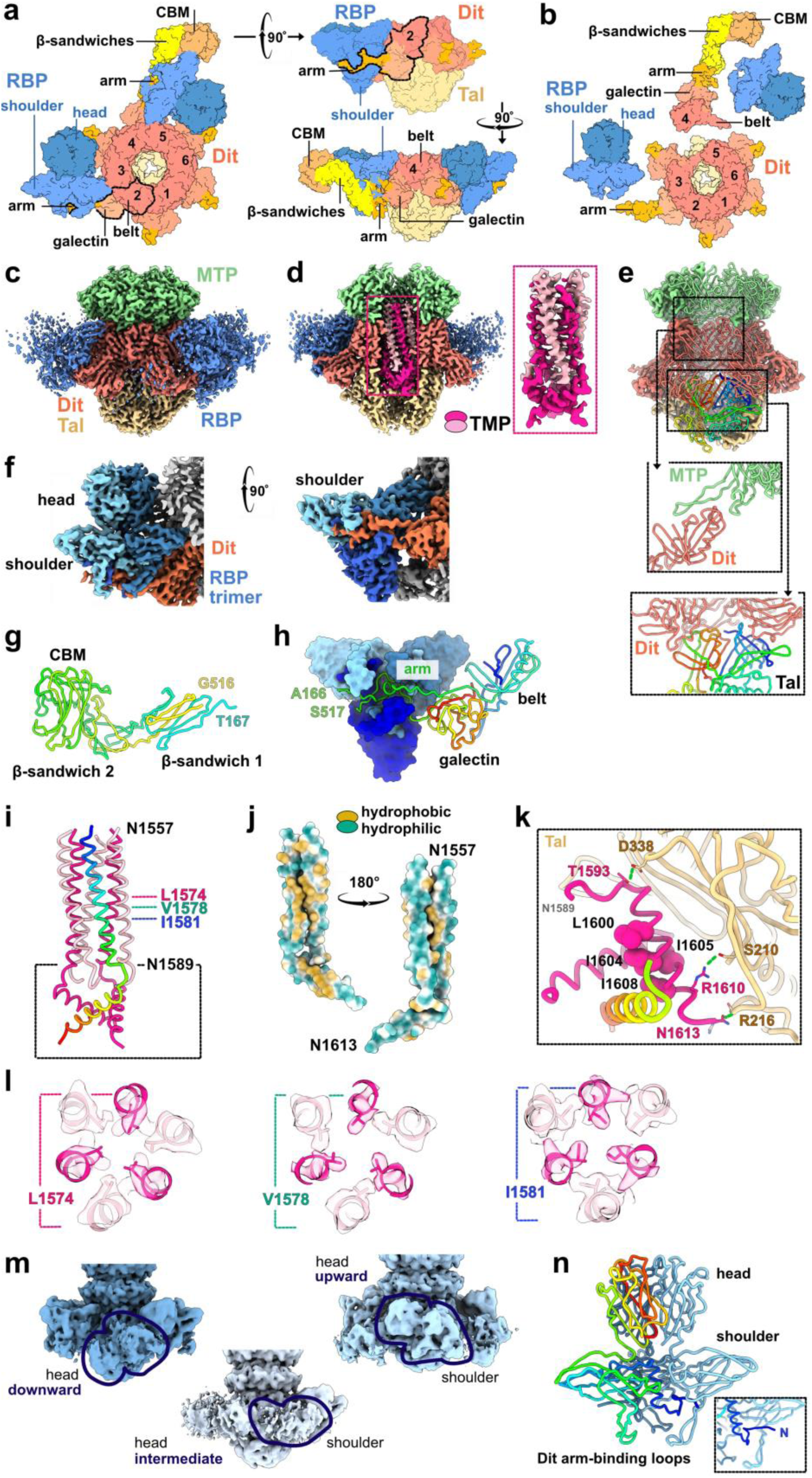
Adhesion device and TMP structures. **a.** Orthogonal views of the overall structural organization of the adhesion device combining cryoEM structures of the Tal-Dit and RBP-Dit complexes with the AlphaFold3-predicted structure of full-length Dit in complex with a RBP trimer (see Methods). The Top view is on the left. Dit and RBP domains are labelled in black and blue, respectively, and Dit subunits are numbered. Dotted lines delineate the contours of the Dit belt and galectin domains and the Dit arm. **b.** Same view as in a, but with one Dit subunit extracted from the adhesion device and the RBP trimers displaced from the Dit arms to which they are attached**. c.** Reconstruction of the adhesion device (C1 symmetry). **d.** Cut-away view of reconstruction shown as in a. The pink box shows the TMP densities. **e.** Reconstruction of the adhesion device (C3 symmetry) and ribbon representation of the MTP-Dit-Tal model. The black boxes highlight protein-protein interfaces. One Tal subunit is rainbow-colored. **f**. Orthogonal views of the 3D reconstruction of the RBP-Dit complex, shown in the same orientations as in a. **g.** Rainbow-colored ribbon representation of the AlphaFold3 predicted structure of the Dit extension. **h.** Surface and ribbon representations of the RBP-Dit model shown in the same orientation as the right view in f. **i.** Ribbon representation of the TMP model, with one subunit rainbow-colored. **j.** Surface representation of a long TMP subunit highlighting hydrophobic and hydrophilic regions within and at the surface of the bundle, respectively. **k.** Close-up of the TMP C-terminal helices. Hydrophobic residues buried within the three-helix bundle are shown as spheres and labelled in black. TMP residues (pink labels) and Tal residues (pale yellow labels) involved in hydrogen bonds (green dotted lines) are shown as sticks. **l.** Reconstruction and ribbon representations of the TMP 6-helix bundle, highlighting focused L1574, V1578 and I1581 as sticks. **m**. 3D classes showing different RBP trimer positions. **n.** Ribbon representation of the RBP trimer with one subunit rainbow-colored. The inset highlights domain swapping between the shoulder domains.

For atomic model building, component-specific reconstructions were performed with symmetry imposed (Supplementary Table 1 and Fig. 2, 5): C6 for the Dit hexamer (2.6 Å overall resolution) and C3 for the Tal trimer (2.7 Å overall resolution). The C3-symmetrized reconstruction was also used to model the TMP structure and to analyse protein-protein interfaces involving the MTP, Dit, Tal and TMP (Fig. 4e,k). Localized 3D refinement focused on a single RBP trimer produced a reconstruction at 3.5 Å overall resolution in which we modeled a RBP trimer bound to a Dit subunit (Fig. 4f; Supplementary Table 1 and Fig. 2, 5, 6).

**Fig. 5.**
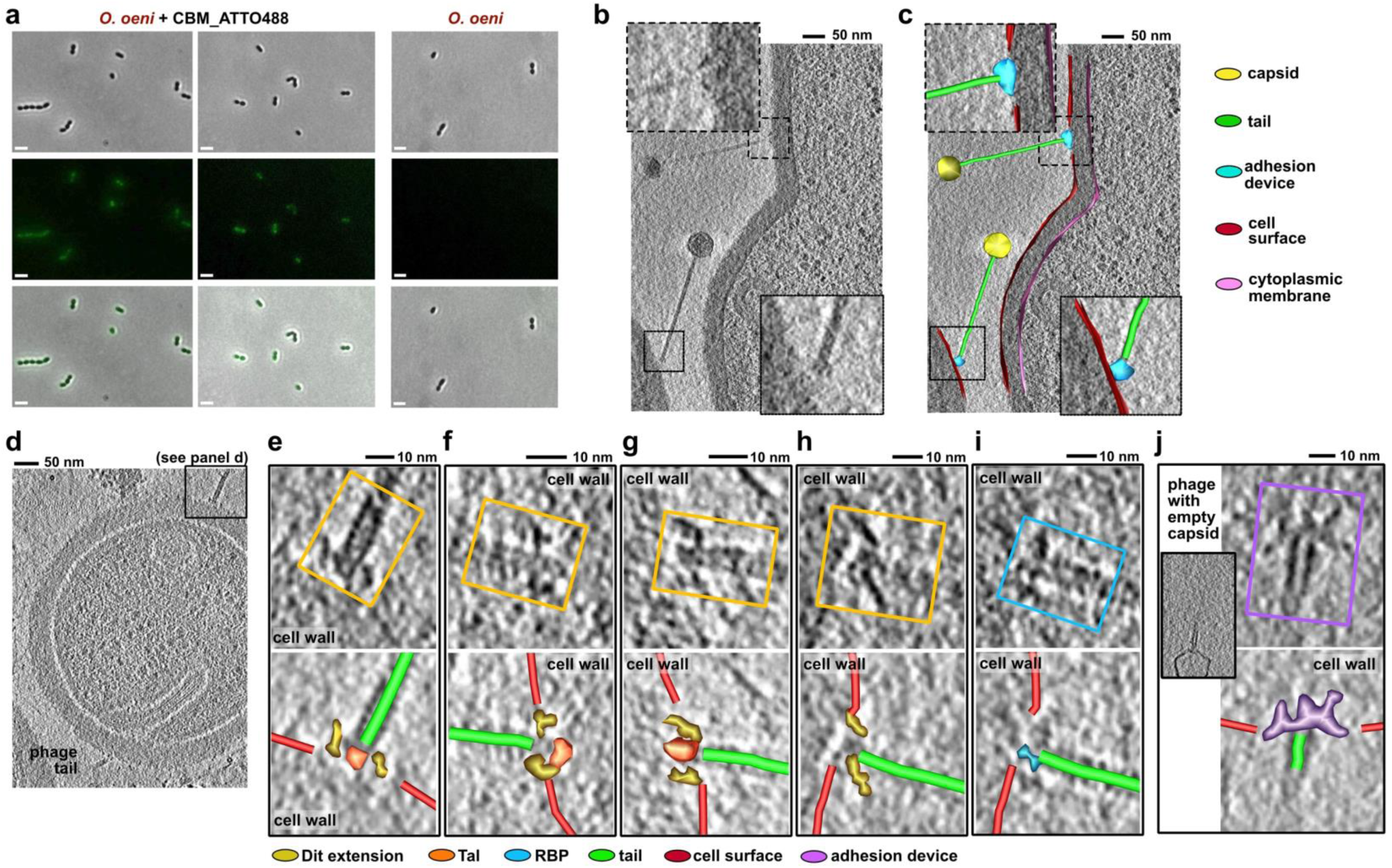
OE33PA interactions with its Gram-positive host. **a.** Fluorescence microscopy images showing the OE33PA Dit’s CBM binding to the *O. oeni* host cell surface. Top row: contrast images; Middle row: fluorescence images; bottom row: overlay between contrast phase and fluorescence images. Scale bar: 2 µm. No fluorescence is detected in the absence of the protein. **b-c.** Z-sections of a tomogram showing OE33PA phage particles interacting with the *O. oeni* cell surface (enlarged views in the insets). Panel c shows the segmentation overlay, highlighting the different components of the phage particles and the different conformations of the adhesion devices **d**. Z-section of a tomogram showing a membrane invagination and a phage tail tip bound to the cell surface. **e-i.** Z-sections of tomograms showing different conformations of the adhesion devices of phages before DNA ejection. The tail tip and adhesion device are highlighted by colored boxes in the top views (Dit extensions in yellow boxes and a RBP trimer in the blue box). Segmentation overlays (contours on successive tomographic slices guided by local contrast differences between the components and their surroundings) are shown in the bottom views. **j.** Z-section of a tomogram showing the adhesion device of a phage after DNA ejection. For panels **b-j**, sections are shown from Z-smoothed versions of the tomograms to improve visualization (see Methods). Scale bars and color codes are indicated in the respective panels. Supplementary Movies, showing the original and Z-smoothed tomograms, with and without segmentation overlays, are provided. These movies improve visualization and allow structural features to be followed across consecutive Z-sections.

The OE33PA Dit is a 659-residue protein comprising a canonical N-terminal belt domain (residues 1-130), which adopts an MTP-like fold and assembles into a hexameric ring, a galectin domain (residues 131-146, 532-659), extended linkers (residues 131-166, 517-535) and a long 3-domain extension that remains unresolved in our 3D reconstructions (residues 167-516) (Fig. 4g ; Supplementary Fig. 9). The extended linkers originating from both the belt and galectin domains project outward to form an arm-like protrusion that engages the N-terminal shoulder domains of the RBP trimers (Fig. 4f-g). The belt and galectin domains closely resemble those of the *L. lactis* phage p2 and the *B. subtilis* phage SPP1 (Supplementary Table 2 and Fig. 7). Loops located on the upper surface of the Dit ring interact with the MTP β3-β6 loop, thereby anchoring the adhesion device to the tail tube (Fig. 4e).

Structure prediction of the full-length Dit using AlphaFold3 yielded a high-confidence model showing that the non-resolved region comprises two β-sandwich domains (residues 168-282, 457-516) followed by a CBM (residues 283-456) (Fig. 4g; Supplementary Table 2 and Fig. 9). These three consecutive domains adopt an elongated arrangement connected to the Dit core via flexible linkers (Fig. 4g; Supplementary Fig. 9), suggesting a mobile extension capable of sampling multiple conformations that are not resolved in our reconstructions (Fig. 4c).

The OE33PA Tal structure comprises four domains and adopts an architecture similar to that of several siphophages, including the *L. lactis* phage p2 (Supplementary Table 2 and Fig. 7, S10). The N-terminal and C-terminal β-sandwich domains of each Tal subunit engage two Dit subunits, thereby accommodating the C6-C3 symmetry mismatch (Fig. 4e). Together, these domains from the three Tal subunits extend the MTP-Dit tube. The dome-shaped conformation of the Tal trimer is stabilized by inter-subunit interactions involving the second and third α/β domains, which seal the central tail channel (Fig. 4c-e). This Tal conformation is consistent with a pre-host-binding conformation of the phage.

Helical densities along the Dit-Tal central channel reveal the hexameric organization of the OE33PA TMP (Fig. 4d; Supplementary Fig. 3, 5, 10). Starting from the density inside the Tal trimer, we modeled the 25 C-terminal residues for three alternating TMP subunits, comprising a short linker (N1589-P1594) followed by an amphipathic α-helix (G1595-N1613) (Fig. 4i,k). Each C-terminal helix forms an approximately 45° angle relative to the tail axis, with its C-terminus located in the center of a Tal subunit and its N-terminus located at the top of the same Tal subunit (Fig. 4i-k; Supplementary Fig. 10). Together, these three helices assemble into a triangular bundle stabilized by hydrophobic interactions (L1600, I1604, I1605, I1608) (Fig. 4j,l). The interface between these three TMP helices and the Tal trimer spans ∼1350 Å² and involves hydrogen bonds (T1593 _TMP_-D338_Tal_, R1610_TMP_-S210_Tal_, and N1613_TMP_-R216_Tal_) (Fig. 4k).

Upstream of the linker, additional 32-residue amphipathic α-helices (N1557-G1588) extend within the central cavity of Dit and part of the upstream MTP ring (Fig. 4d, 4i-j; Supplementary Fig. 10). Furthermore, modeling of the three additional alternating densities inside the Dit-MTP cavity supports an hexameric organization of the TMP (Fig. 4i,l). Although the density in this region is weaker, bulky side chains, including L1574, V1578 and I1581, are distinguishable within the six-helix bundle (Fig. 4l). The interface between this six-helix bundle and the Dit-MTP central channel spans ∼900 Å² (Supplementary Fig. 10). Using different combinations of image processing approaches, including asymmetric refinement, particle subtraction, localized 3D classification, and C3- and C6-symmetric refinement with symmetry relaxation (Supplementary Fig. 3), we confirmed that the C6-to-C3 transition observed in the TMP density is not an artifact arising from imposed symmetry constraints or the contribution of the surrounding hexameric Dit and MTP regions. The difference in the extent of the TMP-Tal and TMP-Dit interfaces may explain why the densities corresponding to the three TMP subunits stabilized by the Tal trimer are better defined than those of the three alternating TMP subunits. Overall, this hexameric-trimeric α-helical assembly at the C-terminal end of TMP appears well suited to accommodate the C6-to-C3 symmetry mismatch within the Dit-Tal complex.

The C1 reconstruction of the adhesion device revealed poorly resolved densities for the RBP trimers, indicating conformational heterogeneity (Fig. 4c,d). To further explore this variability, 3D classification identified three distinct RBP orientations: 1) an upward orientation with the receptor-binding domains, also referred to as the RBP head domains, pointing toward the capsid, 2) a downward orientation with the head domains pointing away from the capsid, and 3) an intermediate orientation (Fig. 4m). Notably, these conformations can coexist within the same adhesion device, with different spatial distributions around the Dit-Tal core (Fig. 4m). The flexible Dit arm loops that engage the RBP shoulder domains appear well suited to accommodate this mobility (Fig. 4f).

We modeled an RBP trimer in the upward orientation bound to a Dit subunit using the localized 3D reconstruction (Fig. 4f, n). The OE33PA RBP comprises an N-terminal shoulder domain (residues 2-149), consisting of a β-sandwich formed by four- and three-stranded β-sheets preceded by an α-helix, connected via a 24-residue loop to a C-terminal head domain (residues 174-261), which also adopts a β-sandwich fold (Fig. 4n). Trimerization of the shoulder domains is mediated by domain swapping within the three-stranded β-sheet and formation of a central three-helix bundle (Fig. 4n). At the base of the shoulder domain, three long loops delineate the Dit arm-binding site. The structure-based identification of the OE33PA protein AWT48019 as a RBP was first proposed in 2021 based on its AlphaFold2-predicted structure^18^. The cryoEM structure determined in the present study further supports this annotation, while not excluding a structural role for the RBP trimers in the mechanical organization of the overall adhesion device. Structural similarity analysis highlights the modular architecture of this protein. While the OE33PA shoulder domain resembles those of *L. lactis* phages p2 and 1358, its head domain is more similar to those of *L. lactis* phages bIL170 and TP901-1 ^32,33^ (Supplementary Table 2 and Fig. 7). Also, the OE33PA RBP, like that of phage 1358, lacks the neck domain present between the shoulder and head domains in phage p2. These similarities between the OE33PA head domain and those of bIL170 and TP901-1 suggest that the OE33PA receptor-binding site is located at the interface between adjacent RBP subunits ^32,33^.

### The Dit CBM is a host-binding domain

The OE33PA CBM is located at the tip of a highly mobile Dit extension, potentially positioning it for initial contact with the host cell. This question is particularly relevant in light of previous findings for *Lactobacillus* phage J-1, whose Dit protein contains two CBM insertions, only one of which was shown to bind to host cells ^34^. To determine whether the CBM can bind to host cells, we recombinantly expressed the protein, labelled it with ATTO-488, and incubated it with live *O. oeni* strain IOEBS277 cells. After washing, samples were mounted on agarose pads and analysed by fluorescence microscopy. Fluorescence imaging revealed decoration of the *O. oeni* cell surface by the labelled CBM (Fig. 5a), supporting its role as a host cell-binding domain. Binding was also assessed using the non-host bacterium *S. thermophilus*, but no detectable fluorescence was observed under the same assay conditions (Supplementary Fig. 11a).

### CryoET of OE33PA bound to *O. oeni*

To obtain structural insights into the host-binding mechanism *in situ*, we performed cryoET on FIB-milled lamellae of cell-bound phages. This approach enabled visualization of the OE33PA adhesion device under native conditions and captured conformational states associated with host binding (Fig. 5b-j and Supplementary Fig. 11b-i).

The cytoplasmic membrane of *O. oeni* is surrounded by an approximately 50 nm-thick polysaccharide-rich cell wall, together forming a typical LAB cell envelope ^35,36^ (Fig. 5b,c and Supplementary Movies 1-5). In several tomograms, we observed intracytoplasmic membrane invaginations, consistent with phage infection, as previously reported in phage-infected *Lactobacillus plantarum* ^36^ (Fig. 5d and Supplementary Movie 6).

Analysis of adhesion devices that were clearly discernible in the tomograms and consistently observed across different tomogram post-processing procedures (Supplementary Fig. 11) revealed multiple conformations of this multiprotein complex, supporting its structural plasticity during host interaction. Notably, we observed the Dit extension in contact with the cell surface (Fig. 5e-g), consistent with a direct role in host binding. The Dit extension, which protrudes from the adhesion device, adopted an elongated conformation, consistent with an extended arrangement of its two β-sandwich domains and the CBM, with the distal tip reaching the cell surface (Fig. 5e,g,h and Supplementary Movies 7-21). More compact, arched conformations of the Dit extension were also observed (Fig. 5e,f and Supplementary Movies 7-16), potentially enabled by flexibility at the junctions between β-sandwich 1, β-sandwich 2, and the CBM. We also observed a rod-like conformation of the Dit extension that did not contact the cell surface (Fig. 5h and Supplementary Movies 22-26), closed conformations of the Tal (Fig. 5e-g and Supplementary Movies 7-21), and a downward-oriented RBP (Fig. 5i and Supplementary Movies 22-26).

Finally, a phage with an empty capsid and an empty tail, indicative of post-DNA ejection, displayed an adhesion device with radial protrusions likely corresponding to Dit extensions (Fig. 5j and Supplementary Movies 27-31). This conformation may represent a state in which the adhesion device flattens against the cell surface upon host binding. Given the limited number of post-DNA-ejection particles in which the adhesion device could be resolved, we cannot determine whether this conformation represents a common intermediate during cell binding and penetration.

## Discussion

The cryoEM structure of the *O. oeni*-infecting phage OE33PA provides the first high-resolution structure of LAB phages and reveals how structural variability shapes phage architecture and host interaction. Importantly, our results present two layers of variability: the organization of the TMP, highlighting structural diversity across phages, and the conformational dynamics of the adhesion device.

TMP are central to phage biology, with roles that include tail length determination ^37–39^, virion stability ^40^, genome delivery ^41,42^ and, in some phages, enzymatic activity ^41,43–45^. While a trimeric organization has long been considered canonical for siphophages ^4,5,11,12,46–48^, our data challenge this view. Specifically, we observed that the C-terminal region of the OE33PA TMP adopts a hexameric arrangement. Consistent with this observation, a hexameric organization has also been reported for the TMPs of the myophages P7-1^49^ and PhiTE^50^, albeit in their N-terminal and central regions. Among siphoviruses, the archaeal dsDNA virus HFTV1 provides a unique and compelling example of a hexameric TMP resolved in density from the N- to the C-terminus ^51^. At the N-terminal end, the TMP interacts with a TMP-DNA spacer, whereas at the opposite end, three TMP subunits form helices that interact with the Tal trimer, while the remaining three TMP subunits bend backward to interact with the Dit subunits (Supplementary Fig. 12).

These findings raise questions regarding TMP stoichiometry: does it vary among phages, or has a hexameric arrangement been overlooked in previous structures? To address this, we re-examined available 3D reconstructions and corresponding models of several phage TMPs (Supplementary Fig. 12a). For the siphophages DT57C ^47^and 80α ^11^, data support trimeric TMP assemblies (Supplementary Fig. 12). Notably, in phage 80α, the N-terminal end of the TMP interacts with the tail completion protein with a 1:3 stoichiometry, consistent with the trimeric assembly observed at its C-terminal end in the adhesion device ^48^. By contrast, analysis of the adhesion device of the *Mycobacterium smegmatis*-infecting siphophage Douge revealed three previously unassigned densities within the central channel, compatible with three ∼20-residue-long α-helices intercalated between three modelled long vertical α-helices (Supplementary Fig. 12b,c). These additional α-helices are shorter than the three modelled ones, and altogether they form an overall TMP arrangement reminiscent of that observed in OE33PA. Similarly, in the central channel of the *M. smegmatis* phage Bxb1 adhesion device, we identified six well-defined, previously unassigned α-helical densities ∼10 Å above the N-termini of the three modelled α-helices, allowing the fitting of six 15-residue-long α-helices (Supplementary Fig. 12d,e). Taken together, these observations support the existence of at least two TMP organizations and highlight the need for careful interpretation of densities at phage tail tips.

The presence of both full-length and shorter TMP subunits in OE33PA, as also observed in phage Douge, where the TMP undergoes a C6-to-C3 transition, raises questions about the fate of the C-terminal residues of half of the TMP subunits. Three hypotheses could explain this observation: 1) a sliding of three TMP subunits results in an out-of-register α-helical bundle, 2) the C-terminal segments of a subset of TMP subunits adopt flexible conformations, or 3) the C-terminal regions of three TMP subunits are cleaved. Although the resolution of the six α-helical densities within the OE33PA Dit channel is limited, recognizable bulky side chains assembling an extensive hydrophobic core suggest that the six TMP helices are in register (Fig. 4h,j), making the first hypothesis less likely. The absence of clear density corresponding to the eighteen C-terminal residues of three TMP subunits in the asymmetric (C1) and symmetrized (C3, C6) reconstructions is consistent with the hypothesis that these regions would adopt flexible conformations that could be accommodated within the available space. This interpretation is supported by the cryoEM structure of the hexameric TMP of HFTV1, which contains six full-length TMP subunits. In this structure, the C-terminal regions of three alternating subunits do not form α-helices but instead adopt elongated conformations that bend back towards the capsid and interact stably with Dit. In Bxb1, additional densities extending towards the capsid are also visible for the corresponding regions of alternating TMP subunits at the end of the six-helix bundle, suggesting that these subunits are likewise full-length. These observations make partial cleavage of half of the TMP subunits less likely, although this possibility cannot be definitively excluded based on the available structural data.

Within the analysed region of the OE33PA internal channel, spanning the connector and the connector-tail junction, we observed only density compatible with DNA (Fig. 3f and Supplementary Fig. 8). No structure resembling the Tail Completion Protein (TCP) observed in phage 80α, which bridges the TMP and DNA, was resolved in this region. However, this does not exclude the possibility that OE33PA encodes a TCP that was either not resolved in our reconstruction or is not present in the mature phage particle. We identified protein AWT48011 as a candidate TCP-like protein based on its high-confidence AlphaFold3-predicted structure, which adopts an elongated α/β fold resembling that of the TCP resolved by cryoEM in phage 80α and those of AlphaFold3-predicted TCP structures from other siphophages (Lambda, HK97, SPP1), as reported by Kizziah et al. ^48^ (Fig. 1a and Supplementary Fig. 9). Moreover, AlphaFold3 prediction of this protein in complex with the N-terminal region (residues 1-67) of six TMP subunits suggests interactions similar to those observed in phage 80α ^48^ (Supplementary Fig. 9). As in the TCP of phage 80α, the OE33PA TCP-like protein has a positively charged surface at the opposite extremity that could mediate interactions with the phage genome (Supplementary Fig. 9). One possible scenario is therefore that the OE33PA TCP-like protein interacts with the N-terminal region of the TMP and with the phage genome, as shown for the TCP of phage 80α. Alternatively, consistent with direct interactions between DNA and TMPs reported for the siphophage DT57C ^47^ and the myophage T4 ^52^ the OE33PA TCP-like protein may associate transiently with the tail during assembly, allowing the N-terminal region of the TMP to interact directly with the phage genome in the mature phage particle.

The intrinsic conformational variability of both the RBP trimers and the three-domain Dit extension, observed in OE33PA prior to host engagement, likely enhances the ability of the adhesion device to sample the surrounding environment. Such structural plasticity may facilitate receptor searching and initial host recognition, thereby providing a mechanistic advantage during infection. The movement of the OE33PA RBP and the Dit-RBP interaction resemble features described for the *L. lactis* phages p2 and 1358 ^7,9,53^. In phage p2, X-ray structures of the adhesion device revealed that it can adopt two conformations: a resting state, in which all the RBP head domains have an upward orientation toward the capsid, and an activated state, in which all the RBPs rotate downward positioning their head domains toward the host surface and enabling receptor binding ^7,9^ (Supplementary Fig. 13). In this system, the upward conformation is thought to be stabilised by the Dit arm of a second Dit ring, or by an extension of the MTP ^7^. In contrast, the OE33PA adhesion device lacks both a second Dit ring and an extra MTP domain, leaving the RBP trimers unconstrained and able to adopt upward, downward, and intermediate conformations (Fig. 4k). Notably, AlphaFold3 predicted the structure of a RBP trimer in a downward conformation in a partial OE33PA adhesion device, similar to the activated state of the phage p2 adhesion device (Supplementary Fig. 13). The RBP trimer is anchored to the Dit arm, positioned between the Dit extension and the central ring (Fig. 4 a,b,e,f; Supplementary Fig. 13). We propose that this arrangement may couple host cell recognition by the Dit CBM to changes in RBP orientation. Specifically, CBM binding to the cell surface could transmit mechanical constraints through the Dit extension and arm, promoting rotation of the RBP trimer towards the downward conformation that may favour stable phage attachment. The observation that Dit extensions can be either bound or unbound to the cell surface (Fig. 5e-h) suggests that different binding states may allow the adhesion device to adjust its contacts with the host surface.

Consistent with a role in host recognition, the fluorescent cell-binding assay showed that the OE33PA Dit CBM binds to the surface of *O. oeni*, whereas no detectable binding was observed with *S. thermophilus* (Fig. 5a; Supplementary Fig. 11). This difference may reflect differences in the cell surface composition of these two Gram-positive bacteria. Recent structural characterization of cell wall polysaccharides from *O. oeni* strain IOEBS277, the host of OE33PA, identified two major polysaccharides exposed at the cell surface ^35^. One is a heteropolysaccharide with repeating units containing a branched hexasaccharide and a glycerol residue linked by phosphodiester bonds, whereas the other consists of a (1→6)-linked β-D-galactofuranan chain partially substituted at the C-2 hydroxyl group with β-D-glucopyranose. These polysaccharides are potential candidates for the cell surface receptors recognized by the OE33PA CBM, although their involvement in CBM binding has not been established. The cell wall polysaccharides of *S. thermophilus* are structurally diverse and include rhamnose- and glucose-containing polysaccharides, as well as exopolysaccharides with varying compositions ^54^. These polysaccharides have been identified as receptors for *S. thermophilus* phages P738 ^55^ and 9871 ^56^, respectively. Differences between the reported cell-surface polysaccharides of *O. oeni* and *S. thermophilus* could contribute to the observed difference in CBM binding. However, the nature of the glycan recognized by the OE33PA CBM remains unknown, and further work is needed to identify the cell surface component mediating this interaction.

Host cell attachment is expected to trigger downstream structural rearrangements that ultimately lead to TMP release and DNA ejection. Any structural perturbation that destabilizes the closed conformation of the Tal trimer, which encloses the C-terminal end of the OE33PA TMP (Fig. 4i), could induce TMP release. The amphipathic nature of the OE33PA TMP C-terminal end (Fig. 4h) is consistent with a potential role in destabilizing and/or disrupting the cell envelope. However, the impact of TMP variability, including oligomeric transitions, on the infection process remains to be determined.

Based on the structures of the adhesion devices of the phages p2, which revealed the “articulation” of RBP trimers around the Dit arm ^7^ (Supplementary Fig. 13a), and OE33PA, we sought to identify conserved structural features that may enable this movement and to uncover additional articulated phages. The notion of “articulation” differs from general structural flexibility in that it involves a relatively rigid-body rotation of the RBP trimer around a defined pivot. The primary defining features are the presence of RBP shoulder domains containing Dit-binding loops and a Dit arm domain. A secondary feature is a short Tal protein restricted to structural domain ^6^. We previously reported that two *L. lactis* phages, PhiLC3 and Q33, display these features, and AlphaFold2 structural predictions supported a *bona fide* interaction between their Dit arm and RBP shoulder domains ^57^. Using these criteria, we identified several candidate articulated phages infecting monoderm Firmicutes and Actinobacteria (Supplementary Fig. 14). AlphaFold3-predicted structures of Dit-RBP complexes showed interactions between the Dit arm and RBP shoulder domains similar to those observed in phages OE33PA and p2 (Supplementary Fig. 14). However, no candidate articulated phages were identified among mycobacterium-infecting phages in the actinobacteriophage database. Regarding the potential functional advantage of articulation, we hypothesize that initial interactions mediated by accessible RBP heads of upward-oriented conformation, or by downward-oriented RBPs that also occur in solution, as shown by the OE33PA structures, could transiently retain the phage at the cell surface during the initial stages of host recognition. This initial interaction could allow additional RBPs to engage their receptors and/or promote RBP rotation under the mechanical constraints imposed by cell surface attachment. Such a mechanism could facilitate productive host binding and, consequently, infection, while potentially reducing unproductive interactions with non-host cells.

In summary, our results update two aspects of phage biology: the structural mechanism of adhesion device and the stoichiometry and architecture of the TMP. Rather than being fixed and uniform, both emerge as versatile and adaptable features suited to the host cell surface, thereby documenting the structural and functional diversity of phages.

## Methods

### Phage production

OE33PA was amplified in the *O. oeni* IOEBS277 bacterial strain and purified as described in Chaïb et *al.* ^58^. The final phage pellet was resuspended in 25 µL of 50 mM Tris-HCl pH 7.5, 100 mM NaCl, 8 mM MgSO4. Phage quality was assessed by negative staining EM ^58^.

### CryoEM grid preparation and data collection

3.5 µL of phage sample was applied to a 300-mesh copper Quantifoil R1.2/1.3 holey carbon grid previously glow-discharged for 45 sec at 25 mA using a GloQube system (Quorum Technologies). The grid was blotted for 5 sec and vitrified in liquid ethane using a Vitrobot Mark IV (Thermo Fischer Scientific). CryoEM grids were screened at the Institute for Structural Biology (Grenoble, France) as part of the Structure-tO-Solution (SOS) pipeline ^59^ offered by ESRF under proposal MX-2556. High-resolution data were collected on a Titan Krios (Thermo Fisher Scientific) operated at 300 kV, equipped with a Gatan K3 direct electron detector and Quantum LS imaging filter, on the CM01 beamline of the European Synchrotron Radiation Facility (Grenoble, France; doi.org/10.15151/ESRF-DC-2493362553) ^60^ . Movies were collected in super-resolution mode binned twice at 105,000 x yielding a pixel size of 0.839 Å, with a total dose of 40 e^-^/ Å ^2^ (40 frames/movie) and a defocus range of −0.8 to −2.2 µm.

### CryoEM image processing and single particle analysis

Image processing was performed using single particle analysis in CryoSPARC v4.5.3 ^61^ to obtain 3D reconstructions of the OE33PA’s capsid, connector, tail and adhesion device. The workflows and details of data processing are presented in the Supplementary Fig. 1 and 2. Multi-frame movies were motion-corrected using Patch Motion Correction, and CTF were estimated using Patch CTF. Particles for the capsid and adhesion device were first automatically picked using blob picking, followed by template picking with 2D templates generated from the blob-picked particles.

DNA-containing capsid particles were selected combining 2D classification and ab initio reconstruction of three volumes without imposing symmetry (C1), and subjected to homogeneous refinement (C1). Particles were then 3D classified into four classes with icosahedral symmetry (I) imposed. Particles of the class producing the most resolved 3D reconstruction were subjected to homogeneous refinement (I) to produce the final capsid 3D reconstruction at an overall resolution of 2.8 Å, which was used to model the structure of the capsid asymmetric unit. Based on the procedure described by Hou et al. to calculate 3D reconstructions of the neck ^62^, these particles were then symmetry expanded (I) and a cylindrical mask covering the 5-fold vertex was generated using

ChimeraX^63^ and used for localized 3D classification (C1) to search for the tail. We selected the class with tail density and measured the distance between the capsid center and the capsid-portal junction and used it to re-extract particles centered at the capsid-portal junction. Re-extracted particles were subjected to homogeneous refinement with C5 symmetry imposed to finely align the capsid, and C5 symmetry expanded. The expanded particles were subjected to localized 3D classification using a mask on the connector, and the 3D class showing C12 and C6 symmetries was subjected to localized refinement (C1, mask on the connector). These tail-aligned particles were C12 and C6 symmetry expanded to perform localized refinements with masks on the portal-adaptor and tail-terminator-stopper-MTP regions, and produce final 3D reconstructions at overall resolutions of 2.6 Å and 2.8 Å, respectively. In order to resolve the density present in the connector’s lumen, particles were C12 symmetry expanded and used for a localized 3D classification with a cylindrical mask within the connector. The 3D class showing the most resolved density was subjected to localized refinement with a mask covering the whole connector to produce the final 3D reconstruction at an overall resolution of 2.9 Å. We also re-extracted the particles at the center of the connector (see above) and performed homogeneous refinement with C6 symmetry imposed to finely align the connector. These particles were C6 symmetry expanded and used for a localized 3D classification with a cylindrical mask within the connector. The 3D class showing the most resolved density was subjected to localized refinement with a mask covering the whole connector to produce the final 3D reconstruction at an overall resolution of 3.2 Å.

Adhesion device particles were selected combining 2D classification and ab initio reconstruction of three volumes without imposing symmetry (C1), and subjected to heterogenous refinement (3 classes, C1). Selected particles were subjected to homogeneous refinement without imposing symmetry, imposing C3 symmetry and imposing C6 symmetry. The final 3D reconstructions were obtained at overall resolutions of 3.1 Å (C1), 2.6 Å (C3) and 2.7 Å (C6). The refined particles (C1 symmetry) were then subjected to a global 3D classification to distinguish adhesion devices with different conformations of their RBP, and to localized 3D classifications with masks covering one RBP trimer and the MTP rings (C1 symmetry). The 3D classes showing the most resolved density were subjected to localized refinements with a mask covering the RBP trimer and the MTP rings to produce the final 3D reconstructions at an overall resolution of 3.5 Å and 2.9 Å, respectively. Local resolution estimation was performed for all reconstructions in cryoSPARC (Supplementary Fig. 3 and 4) ^61,64^.

### Model building, refinement, validation and analysis

Structure predictions of the OE33PA virion’s proteins were performed using the AlphaFold3 server (https://alphafoldserver.com/). Predictions were performed by taking into account the stoichiometry of the complexes: hexamer and pentamer of the MCP, dodecamers of the portal and adaptor, hexamers of stopper, tail terminator and MTP, Dit hexamer, and trimers of Tal and RBP. The models were rigid-body fitted into the cryoEM 3D reconstructions with Coot ^65^, and manually refined when necessary. The TMP structural model was constructed de novo into the 3D reconstruction with Coot, starting from the well-defined trimer of C-terminal α-helices within the Tal cavity and going upwards in the Dit tube. The three other helices were built from the previous trimer and rigid-body fitted into the map.

Model refinement (coordinates, occupancy and B-factors) was performed with Phenix ^66^. Secondary structures were restrained, as well as Ramachandran and side-chain rotamers. Phenix was also used to validate the final models and map-model correlations. Structures were analysed using ChimeraX^63^, Coot and PDBePISA^67^.

Representations of the overall structural organization of the adhesion device shown in Fig. 4a,b were generated using ChimeraX ^63^ by combining our cryoEM structures of the Tal-Dit and RBP-Dit complexes with the AlphaFold3-predicted structure of full-length Dit in complex with a RBP trimer. The experimentally-determined and AlphaFold3-predicted RBP-Dit complexes were superimposed onto two different Dit subunits. All components of the adhesion device are represented as surfaces. The models of the hexamric Douge and Bxb1 TMPs were produced using using the “Place Helix Here” function in Coot ^65^ into the experimental cryoEM densities. We generated polyalanine helices solely to assess and illustrate the interpretation of these densities. We then further adjusted the six Bxb1 helices and the three additional Douge helices within their respective densities using the “Fit in Map” function in ChimeraX. The resulting map-model correlation coefficients were 0.82 for Bxb1 and 0.76 for Douge.

### CBM production and labelling

The sequence encoding the OE33PA Dit CBM (residues 283-444), fused to C-terminal hexahistidine (underlined) and ALFA (bold) tags separated by a GGS linker (italics)(<u>HHHHHH</u>*GGS***PSRLEEELRRRLTE**), was inserted into the expression plasmid pET24(+) (Twist Bioscience). This construct was produced in *Escherichia coli* Rosetta 2 (DE3) pLysS cells, grown in Terrific Broth medium, by the addition of 0.1 mM isopropyl β-D-1-thiogalactopyranoside (IPTG) and incubation for 18 hours at 17°C. Cells were harvested by centrifugation (6,000 x g, 15 min, 4°C), resuspended in buffer A (50 mM Tris pH 8, 500 mM NaCl, 10 mM imidazole, 5 % glycerol) supplemented with 1mM phenylmethylsulfonyl fluoride (PMSF) and 0.25 mg/mL lysozyme, and disrupted by sonication. The lysate was cleared by centrifugation (15,000 x g, 30 min, 4°C), and the supernatant was applied onto 1 mL of His•Bind resin^®^ (Merck) for Ni-NTA affinity chromatography. The immobilized proteins were eluted in buffer B (50 mM Tris pH 8, 500 mM NaCl, 5 % glycerol 250 mM imidazole). The CBM was further purified by size exclusion chromatography using a HiLoad^®^ 16/600 Superdex^®^ 75 pg column (Cytiva) equilibrated in buffer C (50 mM HEPES pH 7.5, NaCl 500 mM). The CBM was concentrated to 1.8 mg/mL using Vivaspin^®^ PES centrifugal concentrators, flash-frozen in liquid nitrogen, and stored at −80°C.

### Fluorescence microscopy

The OE33PA Dit CBM was labelled with the Atto-488 NHS ester fluorescent dye (Merck). The CBM (80 µM) was incubated with the dye (1:1 molar ratio) in buffer C for 1 hour at room temperature. Then, the remaining free dye was removed using desalting column, and the labelled protein was centrifuged (13,000 x g, 5 min, 4°C) and dialysed into buffer D (200 mM Sodium Acetate pH4.8, 10 mM CaCl_2_). *O. oeni* and *S. thermophilus* were grown in Man Rogosa and Sharp (MRS) medium at 30°C^58^ and LM17 medium at 42 °C, respectively. 50 µL of *O. oeni* and *S. thermophilus* cells (OD_600_= 0.2-0.4) were diluted into 250 µL of buffer D, centrifuged (5,000 x g, 5 min), and resuspended with the fluorescently-labelled CBM at 400 µg/mL. Cells were incubated with the fluorescent CBM for 20 min at 30°C, harvested and resuspended in 250 µL of buffer D. Live bacterial cells were imaged using agarose pads on a Nikon Eclipse Ti microscope equipped with an Orcaflash 4.0 LT digital camera (Hamamatsu). Exposure times were typically 30 ms for phase contrast and 200 ms for Atto488-CBM. The experiments were performed in triplicate and representative results are shown. Images were analyzed using ImageJ (https://imagej.nih.gov/ij/).

### CryoET sample preparation, data collection, processing and analysis

*O. oeni* cells were grown in 50 mL of MRS medium to an OD_600 nm_ ∼ 0.5, harvested and resuspended in 10 µL of buffer D. OE33PA was purified as described previously to ∼ 10^13^ pfu/mL and diluted (1/10) in buffer E (200 mM Sodium Acetate pH4.8, 10 mM CaCl_2_, 10% glycerol). Cells and phages were incubated for 15 min at 30°C with 10 nm BSA-treated gold fiducial beads (Aurion, cat. No 210.133). 3 µL and 1.5 µL of the cell-phage sample were applied on each side of a 300-mesh copper Quantifoil R2/2 holey carbon grid previously glow-discharged for 45 sec at 25 mA using a GloQube system (Quorum Technologies), and the excess solution was blotted away on one side of the grid before plunge-freezing (Leica EM GP 2).

Cryo-FIB milling was performed using the Aquilos2 cryoFIB/SEM (Thermo Fischer Scientific) at the Pasteur Institute NanoImaging Core facility (Paris, France). Ten grids were screened for optimal ice thickness, and one selected grid was then sputter coated with platinum GIS for 90s. Nineteen lamellae were initially coarsely milled in auto mode using current of 0.1nA to reach the thickness of 800nm. Then, the fine milling was done manually in four steps with the current set to 50pA to reduce lamellae thickness to 150nm. The lamellae were imaged on a 300 kV TEM Titan Krios (Thermo Fischer Scientific) equipped with an energy filter Selectris X (slit width of 10eV), and a direct electron detector (Falcon4i). Tilt-series data were collected using the Tomography (Thermo Fischer Scientific) data collection package, from −50° to +70° (3° increments) using a dose symmetric scheme, resulting in a total electron dose of 120 e^-^/Å^2^, at a magnification yielding a pixel size of 2.453 Å and a nominal −3.5 to −5.0 µm defocus range.

Tilt-series processing was performed using IMOD v5.1.3 and its Etomo interface ^68^. Tilt-series images were aligned using fiducial models, the contrast transfer function (CTF) was estimated using Ctfplotter, and CTF-corrected tomograms were generated ^69^. For structural interpretation, tomograms were cropped to the regions of interest and binned by factors of 4 or 6. The resulting tomograms were then processed using the deconvolution module in IMOD (deconvolution strength, 0.7) and Topaz Denoise 3D (v0.3.18; model, unet-3D, patch size, 256, patch padding, 32) ^70^. Processed tomograms were visualized using the 3dmod interface in IMOD. Individual components of the adhesion device, including the Dit extension, RBP trimer and Tal trimer, as well as the phage tail and the cell surface, were manually segmented in 3dmod using the deconvoluted tomograms. Segmentation was performed by drawing contours on successive tomographic slices, guided by local contrast differences between the components and their surroundings. Contours were delineated across successive slices to reconstruct the three-dimensional shapes of the individual components. The resulting segmentations are presented in Fig. 5. Corresponding views of the tomograms denoised using Topaz Denoise 3D are provided in Supplementary Fig. 11 to demonstrate the consistency of the structural features observed using the two processing approaches. Tomographic sections shown in Fig. 5 are from Z-smoothed versions of the tomograms. The Z-smoothing was performed by applying a centered three- or five-section moving average along the Z-axis to the already reconstructed tomograms. This linear spatial averaging reduces section-to-section intensity fluctuations and uncorrelated background noise while retaining signal that is spatially coherent across several consecutive Z-sections. It therefore improves the visual contrast of the structural features. The smoothing was used solely as a visualization aid and was not required for the detection or interpretation of the structural features. The features shown in the smoothed sections are independently present in the corresponding non-averaged tomograms.

## Acknowledgments

We acknowledge Romain Linares, our local contact for data collection on CM01 at the European Synchrotron Radiation Facility (Grenoble, France). We gratefully acknowledge Thierry Doan for providing access to the fluorescence microscope and Stéphane Tachon at the Pasteur Institute NanoImaging Core facility (Paris, France) for the cryoET experiments.

## Author contributions

Conceptualization: A.G.; Methodology: L.S., A.C., D.P.; Validation: L.S., A.G.; Formal Analysis: L.S., A.G., C.C.; Investigation: L.S., A.C., E.K., D.P., C.C, A.G.; Resources: A.G., A.C., D.P., C.L.M.; Writing - Original Draft: C.C., A.G.; Writing - Review and Editing: A.G., C.C., E.K., D.P., A.C., C.L.M; Visualization, L.S., C.C., A.G.; Supervision, A.G.; Project Administration, A.G.; Funding Acquisition, A.G., C.L.M. All authors read and approved the final manuscript.

## Data availability

The cryoEM 3D reconstructions and the associated structural models have been deposited in the Electron Microscopy Data Bank and Protein Data Bank, respectively, under these accession codes: EMD-58149, PDB 30YS (capsid); EMD-58150, PDB 30YT (capsid-portal); EMD- 58152, PDB 30YV (portal-adaptor); EMD-58153, PDB 30YW (stopper-TT); EMD-58151, PDB 30YU (connector-MTP); EMD-58155 (DNA-portal-adaptor); EMD-58154 (DNA-adaptor-stopper- TT); EMD-58160, PDB 30ZA (tail tube); EMD-58158, PDB 30YY (adhesion device, C6); EMD- 58157, PDB 30YX (adhesion device, C3); EMD-58156 (adhesion device, C1); EMD-58161, EMD- 58162 and EMD-58163 (3D classed of the adhesion device); EMD-58159, PDB 30YZ (Dit-RBP). The tomograms have been deposited in the Electron Microscopy Data Bank, under the accession codes EMD-57915, EMD-57916, EMD-57918, EMD-57919, EMD-57920, EMD-57921 and EMD-57922. Other data are available upon request to the corresponding author.

---

**Supplementary Fig. 1.**
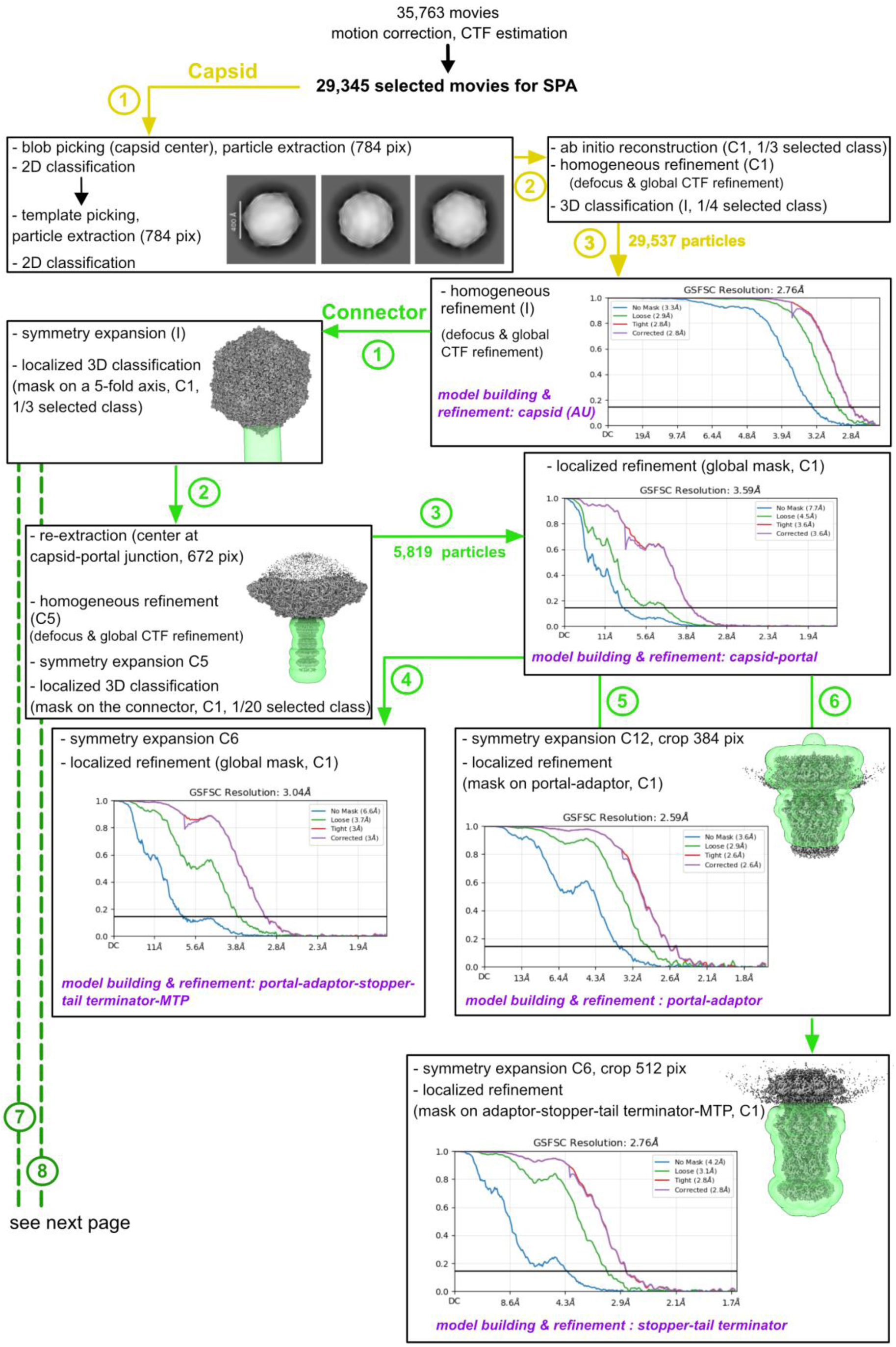
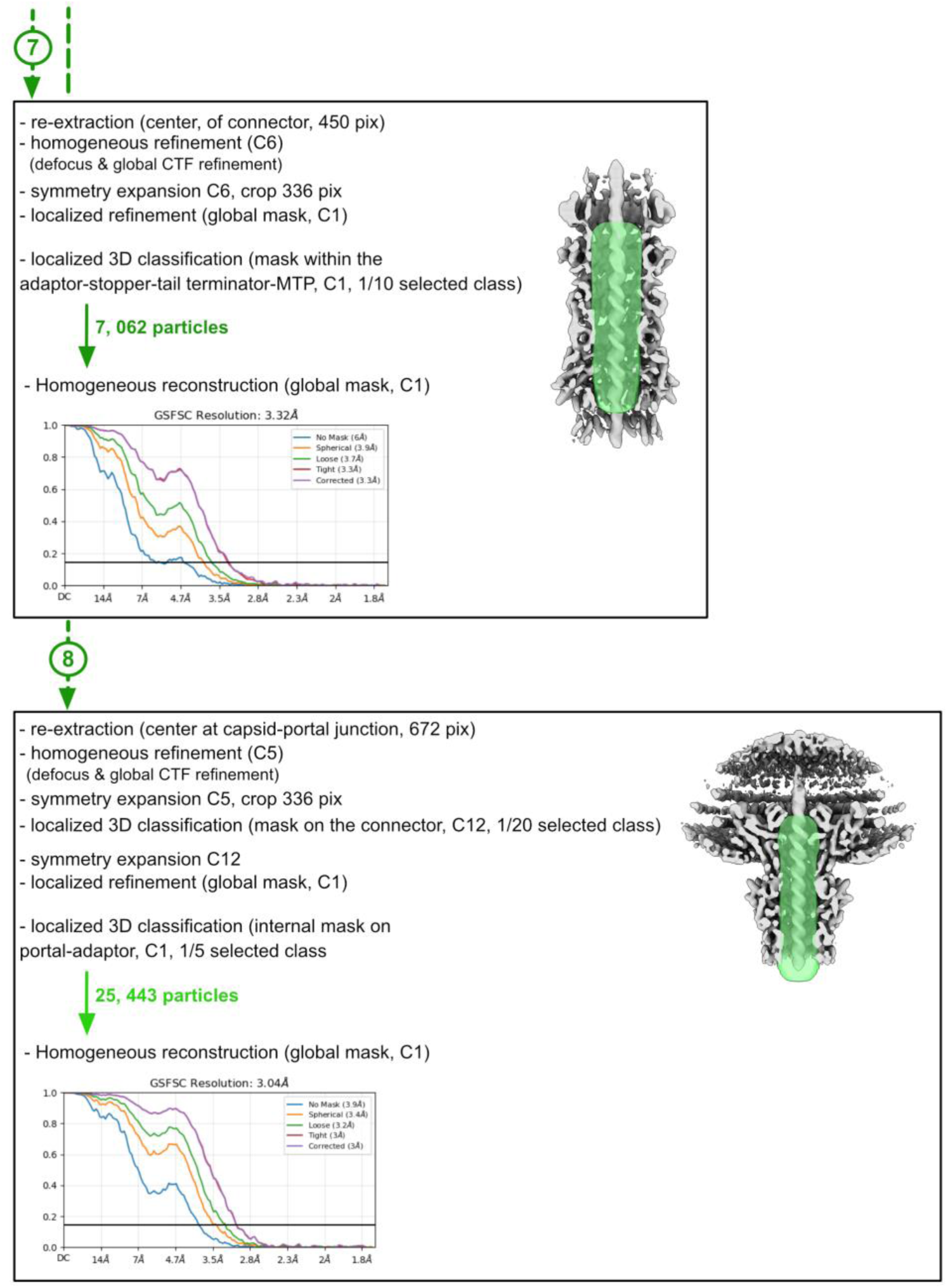
CryoEM structure determination workflow for the capsid and connector. Processing steps of the capsid and connector are shown in yellow and green, respectively. Masks used for 3D classification and localized refinement, and gold-standard FSC curves (cut-off 0.143) are shown. Model building and refinement is indicated in purple.

**Supplementary Fig. 2.**
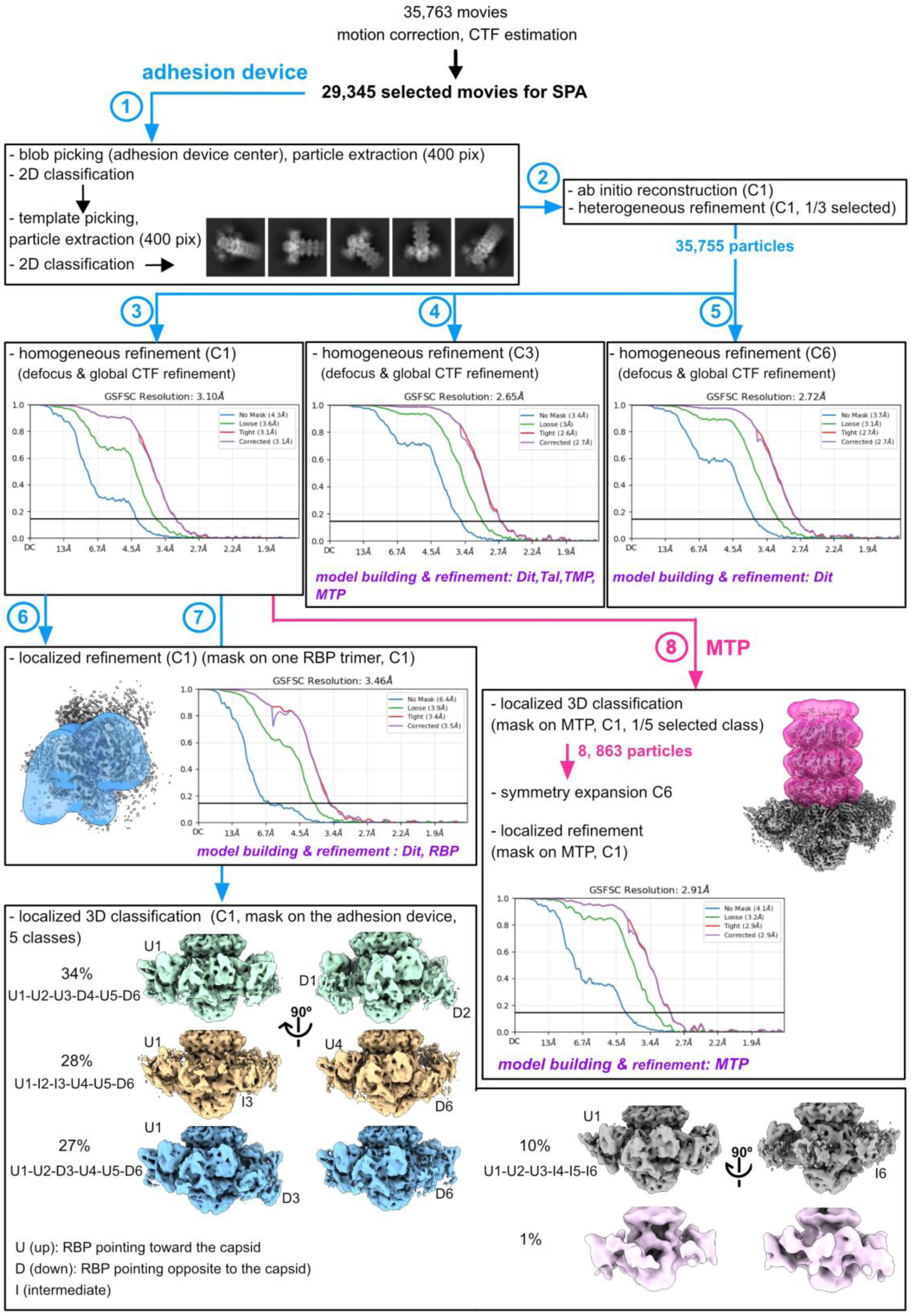
CryoEM structure determination workflow for the adhesion device and MTP. Processing steps for the adhesion device and MTP are shown in blue and pink, respectively. Masks used for 3D classification and localized refinement, and gold-standard FSC curves (cut-off 0.143) are shown. Model building and refinement is indicated in purple.

**Supplementary Fig. 3.**
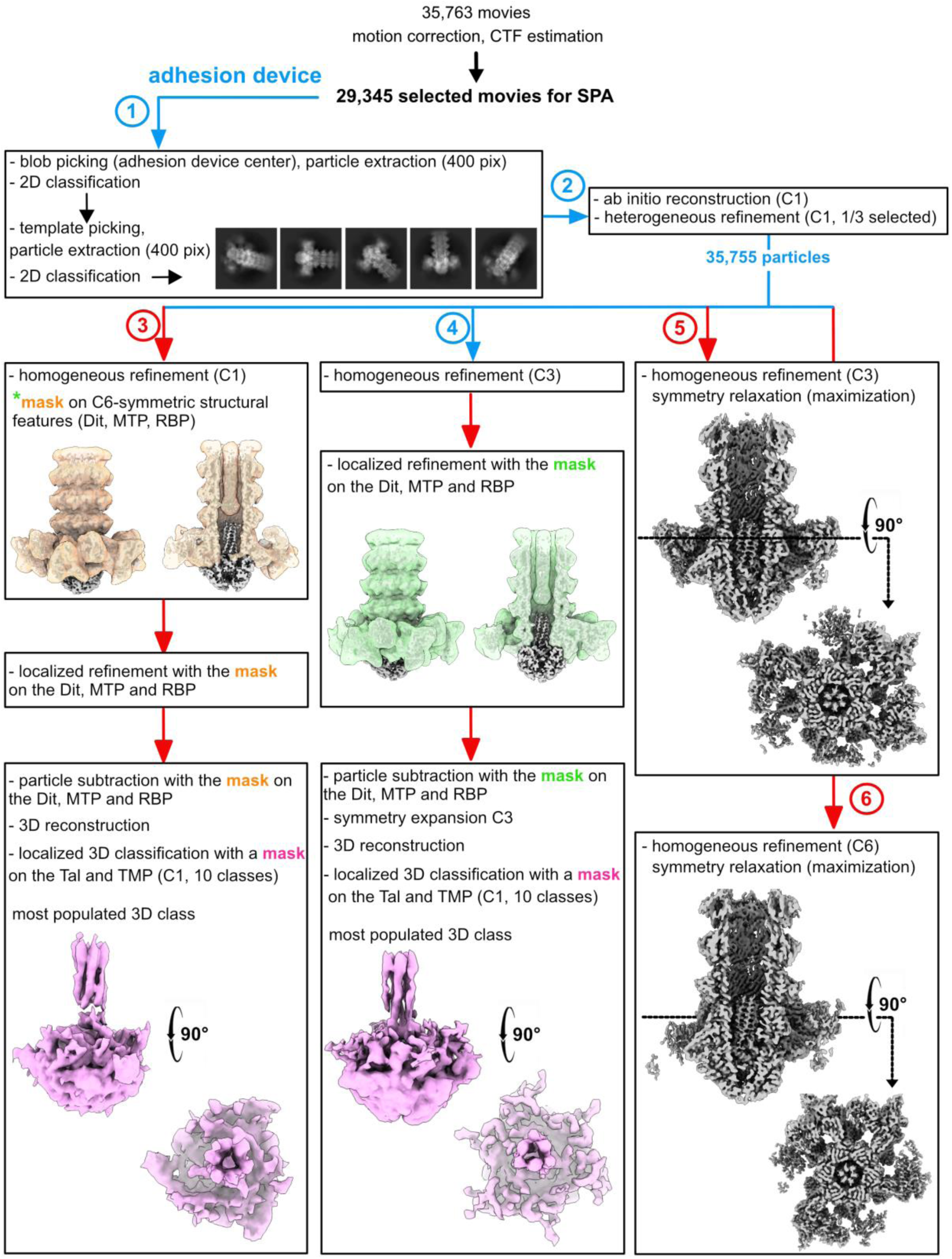
CryoEM structure determination workflow for the TMP. Different image processing strategies, including asymmetric refinement, particle subtraction, localized 3D classification, and C3- and C6-symmetric refinement with symmetry relaxation, consistently confirm the C6-to-C3 transition in the TMP density. Blue arrows refer to image processing described in Supplementary Fig. 2, whereas red arrows indicate additional image processing strategies specifically designed to investigate the TMP structure.

**Supplementary Fig. 4.**
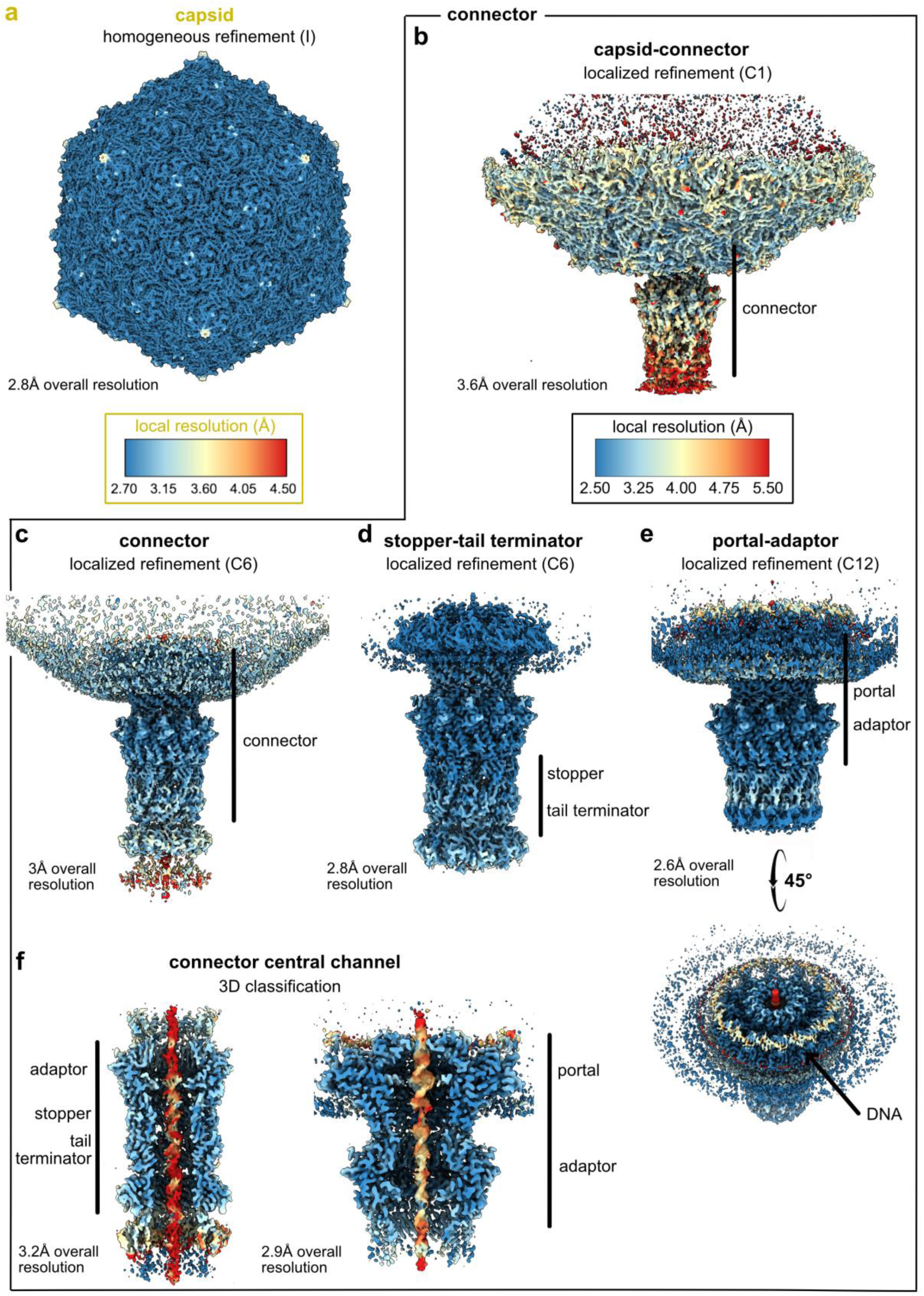
Local resolution distribution of the capsid and connector cryoEM maps. Surface representations colored according to local resolution are shown for the capsid (**a**) and connector (**b-f**). The overall resolution and refinement strategy for each map are indicated in the corresponding panel; the complete information is provided in Supplementary Fig. 1 and Table 1. The minimum value and range of each color scale were selected based on the overall resolution of the corresponding map to highlight meaningful local variations in resolution.

**Supplementary Fig. 5.**
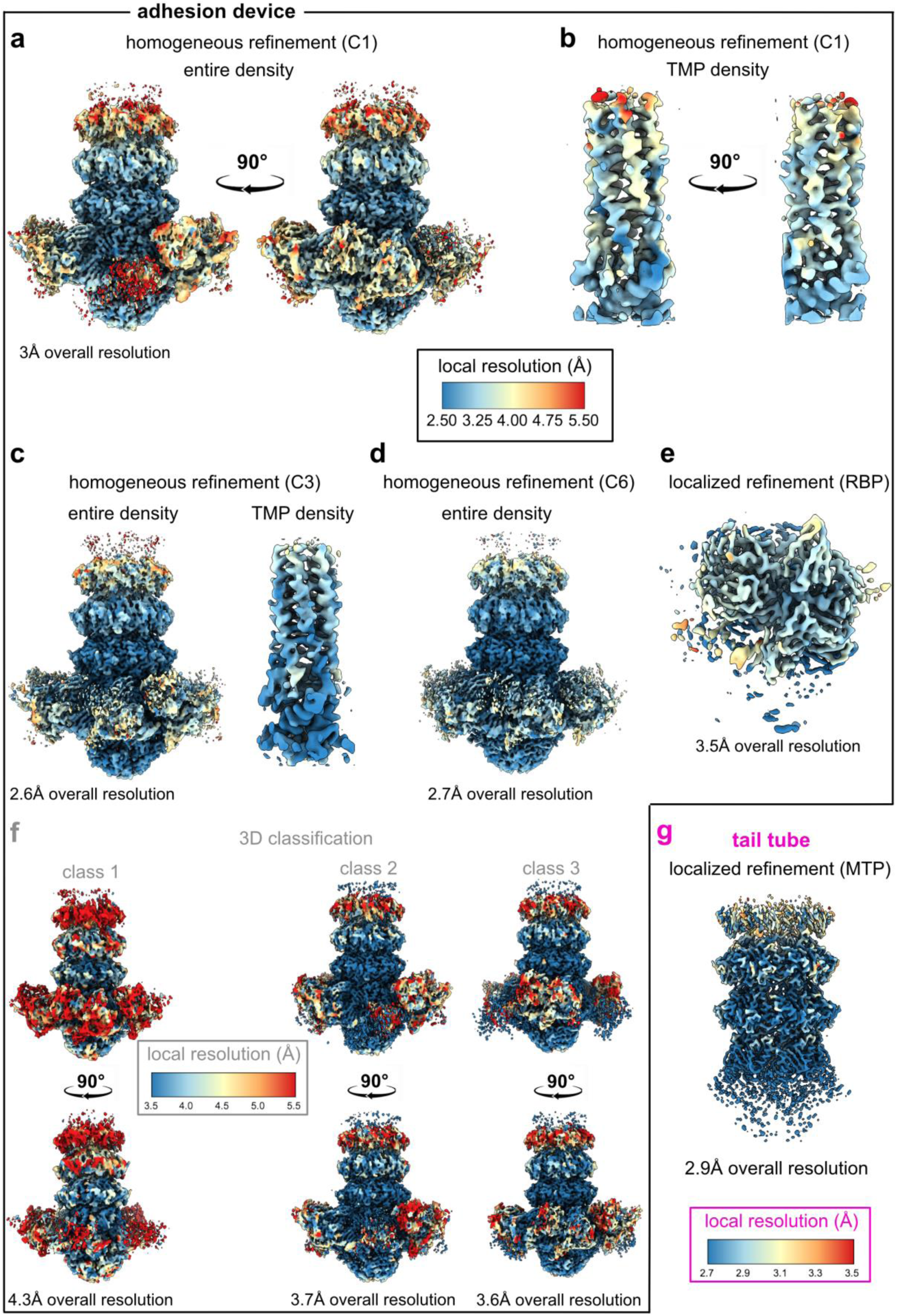
Local resolution distribution of the adhesion device and tail tube cryoEM maps. Surface representations colored according to local resolution are shown for the adhesion device (**a-f**) and tail tube (**g**). The overall resolution and refinement strategy for each map are indicated in the corresponding panel; the complete information is provided in Supplementary 2 and Table 1. The minimum value and range of each color scale were selected based on the overall resolution of the corresponding map to highlight meaningful local variations in resolution. In panels **b** and **c**, only the density corresponding to the TMP is shown, as extracted from the reconstruction of the entire adhesion device.

**Supplementary Fig. 6.**
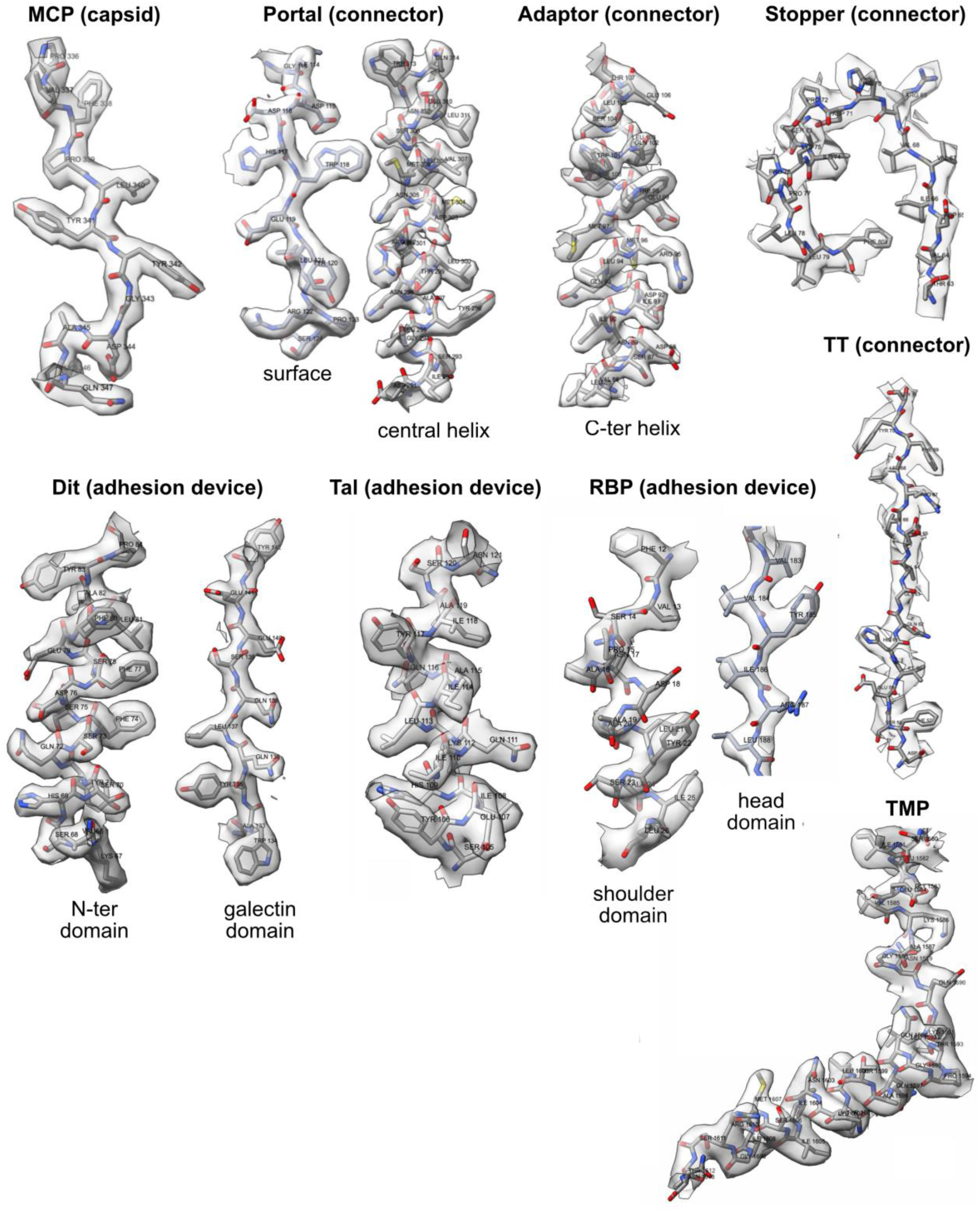
3D reconstructions and model building. Parts of 3D reconstructions and their atomic models are shown as surfaces and sticks, respectively, for the capsid, connector and adhesion device.

**Supplementary Fig. 7.**
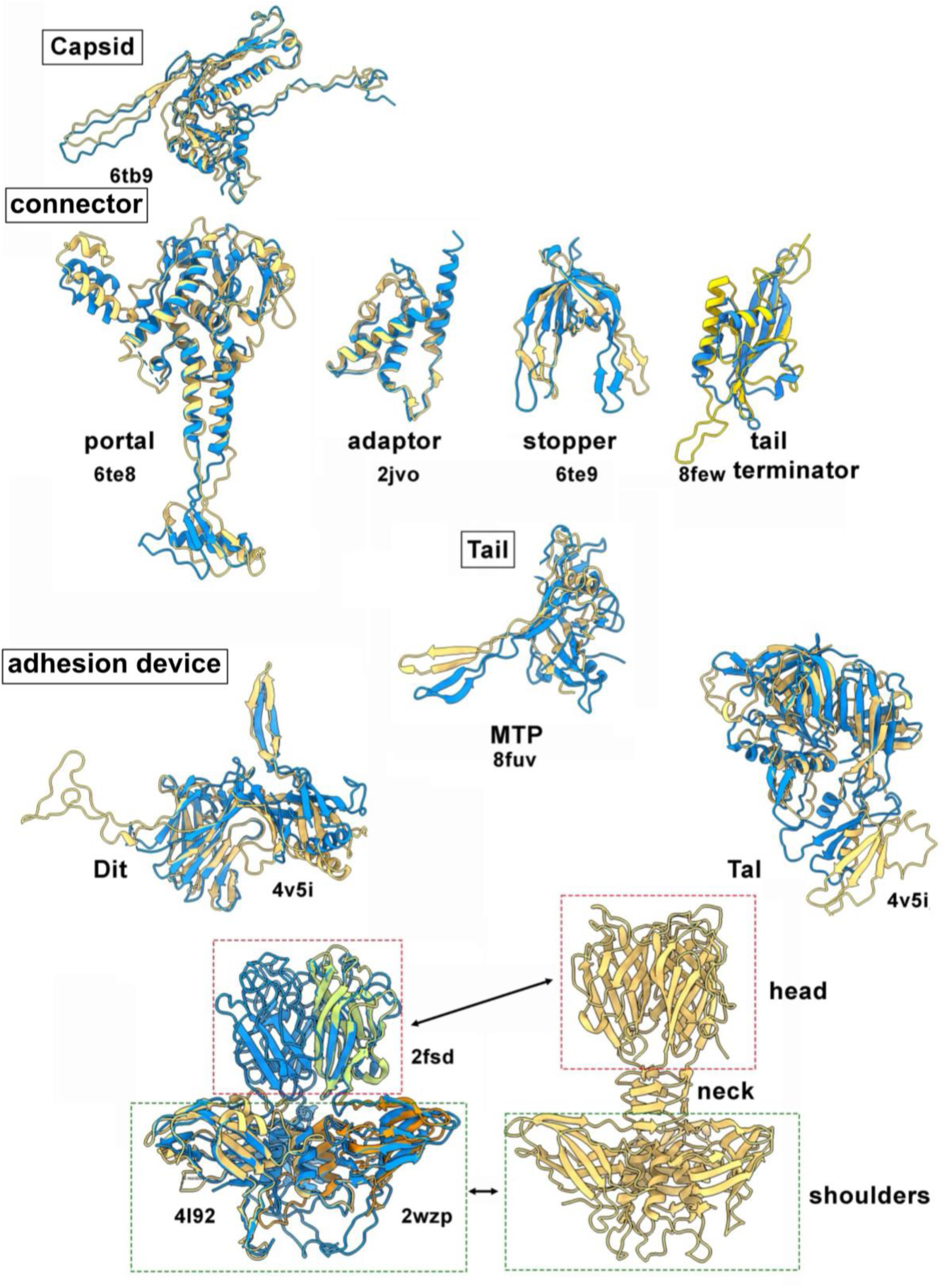
Comparisons between the structures of OE33PA virion components and. **PDB deposited structures.** The structures of the OE33PA virion component are shown in blue and their structural homologs identified by Foldseek are shown in yellow, orange and green. The PBD code is given for structural homologs

**Supplementary Fig. 8.**
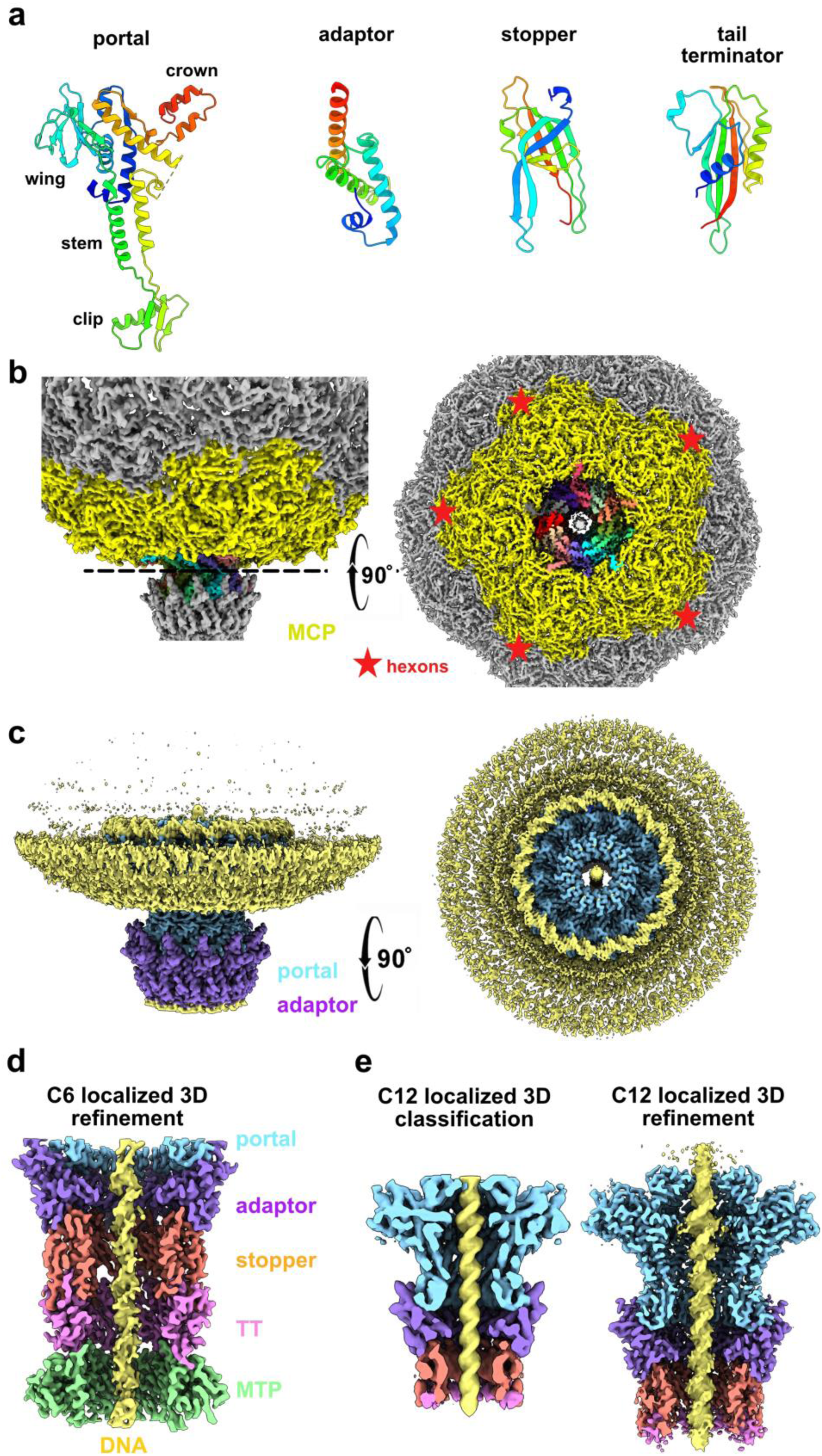
Connector-capsid and Connector-DNA assemblies. **a.** Rainbow-colored ribbon representations of the different connector proteins. **b.** Surface representations of the capsid-portal assembly (cryoEM reconstruction without imposing symmetry). The capsid is yellow and the 12 portal subunits are numbered and shown in different colors. The red stars indicate the five surrounding MCP hexons. **c.** Surface representation of the cryoEM 3D reconstruction of the capsid- portal assembly with C12 symmetry imposed. **d.** Surface representation of the localized 3D reconstruction of the connector with C6 symmetry imposed. **e.** Surface representations of the connector centered at the capsid-portal junction with C12 symmetry imposed after localized 3D classification and refinement.

**Supplementary Fig. 9.**
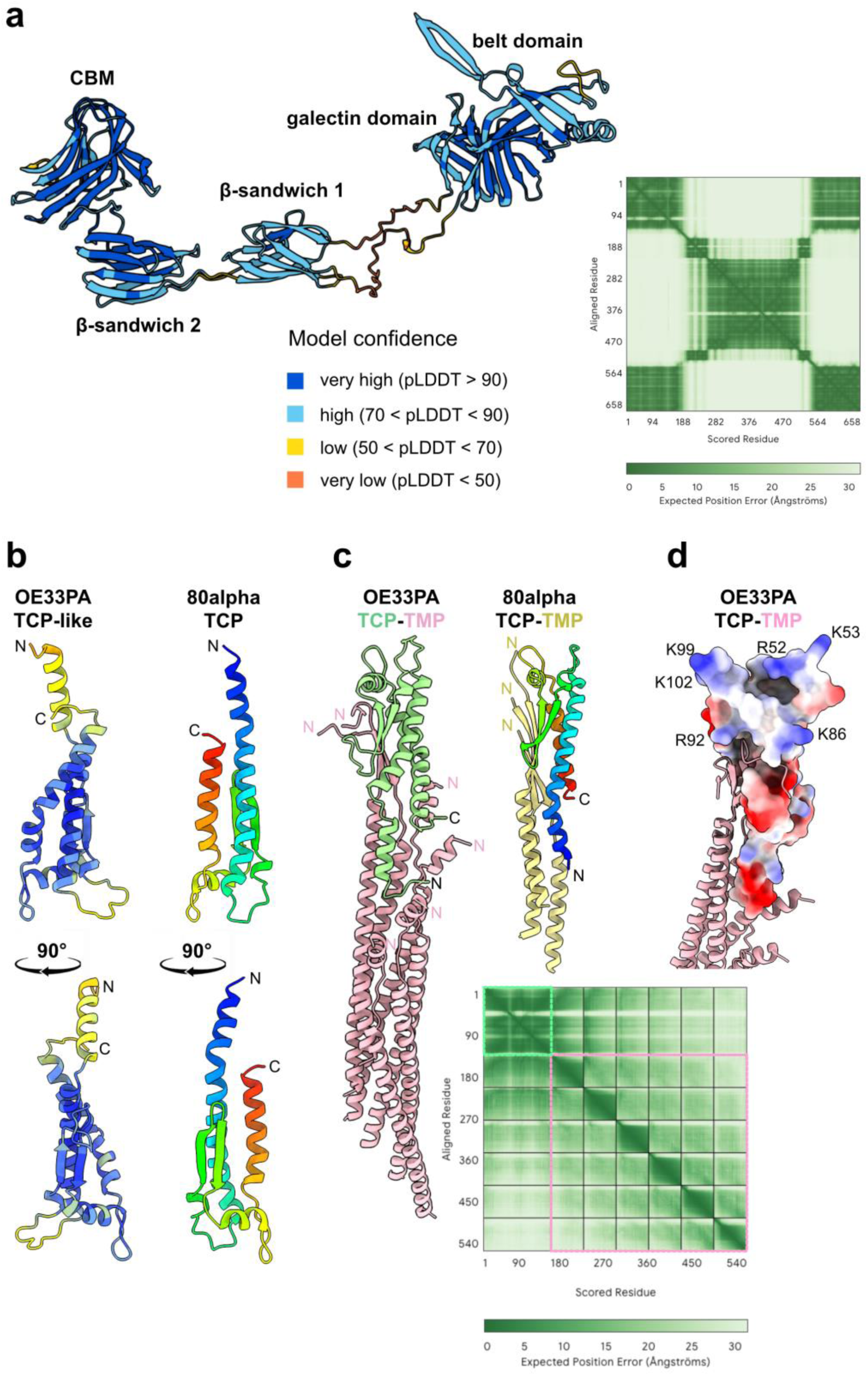
AlphaFold3-predicted structure. **a.** The OE33PA Dit structure is shown as ribbons colored according to pLDDT values. The PAE plot is shown. **b.** The OE33PA TCP-like structure, shown as ribbons colored according to pLDDT values, is compared with the cryoEM structure of the TCP from phage 80α, shown in rainbow colors. **c.** Ribbon representations of the predicted complex between the OE33PA TCP-like protein and six TMP subunits, together with the PAE plot, are shown alongside the structure of the phage 80α TCP-TMP complex for comparison. **d.** The positively-charged extremity of the OE33PA TCP-like protein could potentially interact with the phage genome, as observed for the TCP of phage 80α.

**Supplementary Fig. 10.**
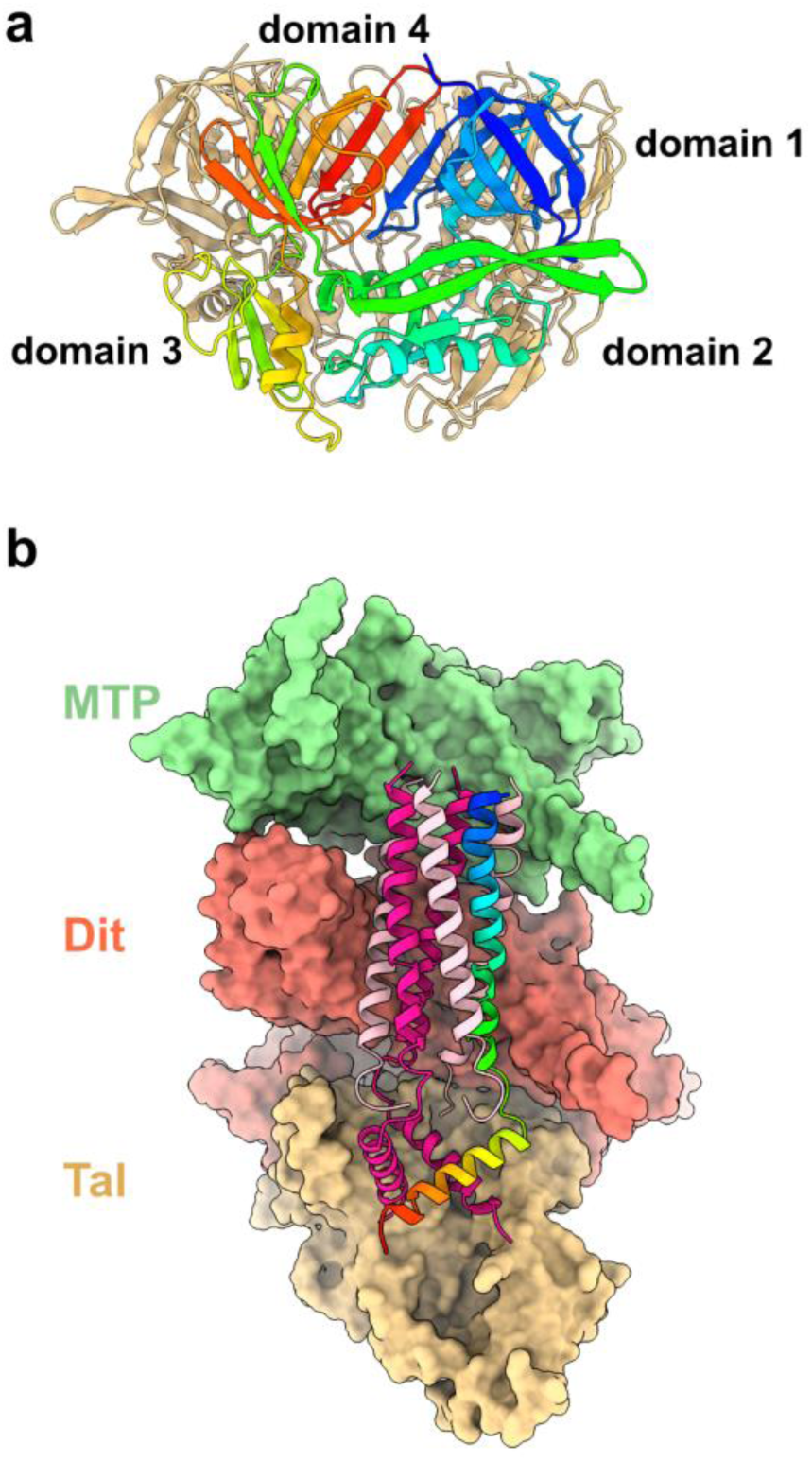
Adhesion device and TMP. **a.** Rainbow-colored ribbon representation of the OE33PA Tal trimer. **b.** Surface representation of the MTP-Dit-Tal assembly with the TMP shown as ribbons. Some subunits of the MTP, Dit and Tal multimers are not shown for clarity. One TMP subunit is rainbow-colored.

**Supplementary Fig. 11.**
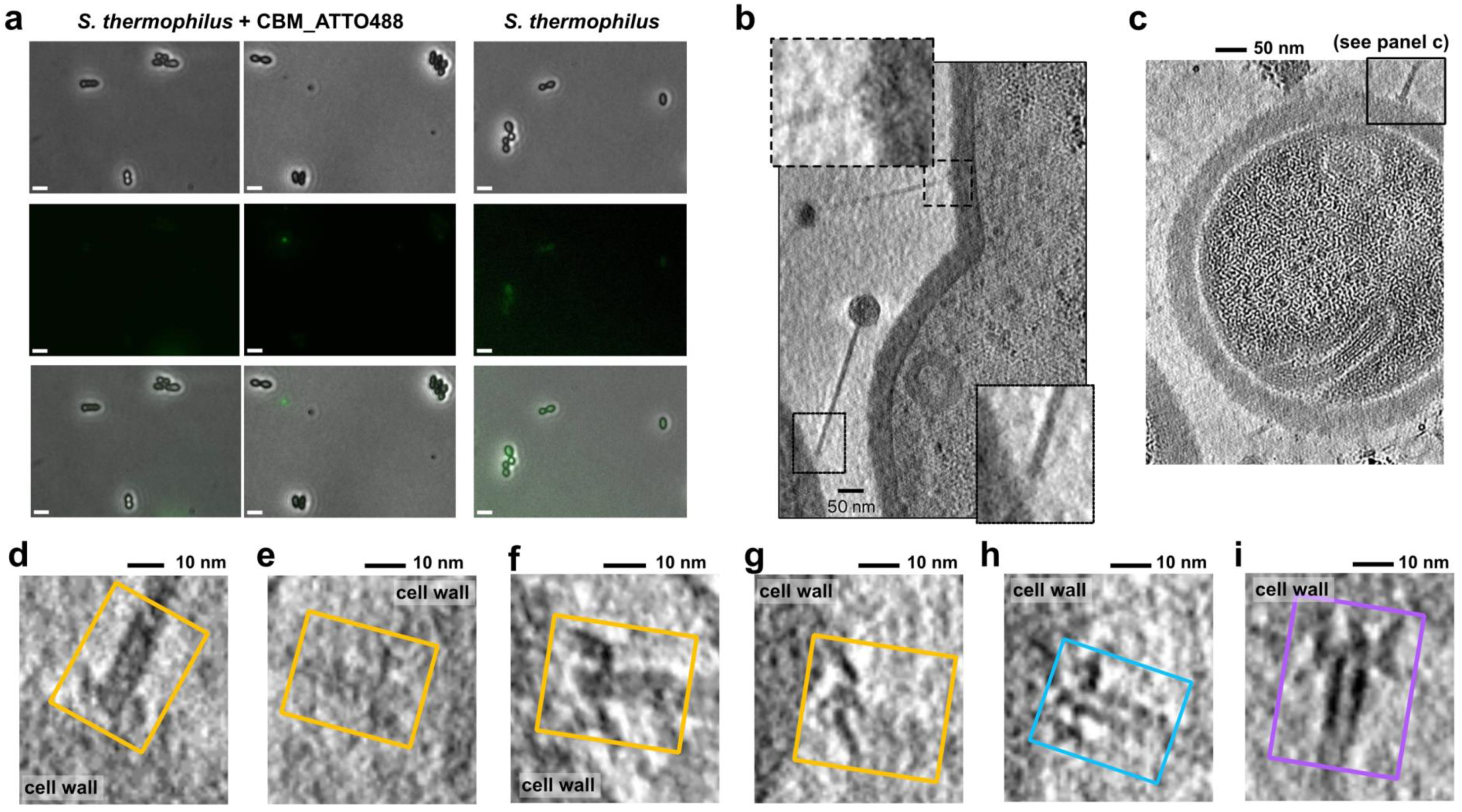
OE33PA interactions with its Gram-positive host. **a.** Fluorescence microscopy images showing that the OE33PA Dit’s CBM does not bind to *S. thermophilus* cells. Top raw: contrast phase images; Middle row: fluorescence images; bottom raw: overlay between contrast phase and fluorescence images. Scale bar: 2 µm. **b-i.** These panels are presented for comparison with those in Fig. 5 to demonstrate the consistency of the structural features observed using two different processing approaches. The panels shown here are sections of tomograms denoised using Topaz Denoise 3D, whereas those in Fig. 5 are sections of tomograms processed using the deconvolution module in IMOD. **b.** Z-section of a tomogram showing OE33PA phage particles interacting with the *O. oeni* cell surface. Enlarged views of different conformations of the adhesion devices are shown in the insets. **c.** Z-section of a tomogram showing a membrane invagination and a phage tail tip bound to the cell surface. The black box indicates the adhesion device shown in panel d. **d-h.** Adhesion devices of phages before DNA ejection, showing different conformations of the Dit extension (d-g) and a downward-oriented RBP (h). **i.** Adhesion device of a phage after DNA ejection (empty capsid), showing extended radial protrusions that likely correspond to Dit extensions. All these structural features can be compared with those shown in Fig. 5.

**Supplementary Fig. 12.**
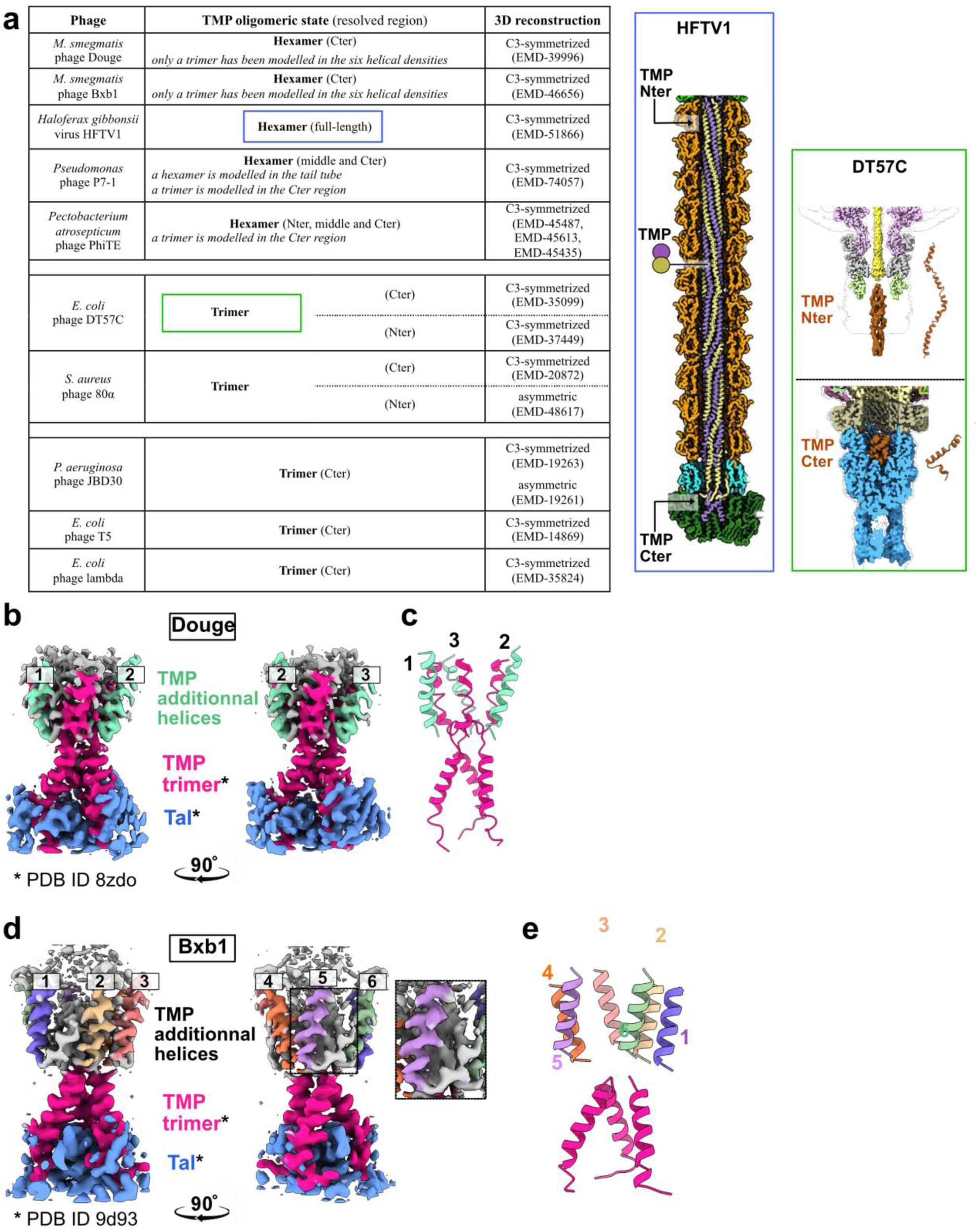
Structural analysis of TMP from various phages. **a.** Summary of the structural evidence supporting the assignment of TMP oligomeric states. Colored boxes show images of the HFTV1 and DT57C TMP assemblies adapted from the original publications ^1,2^. **b.** 3D reconstruction showing the density corresponding to the Douge TMP. **c.** Ribbon representation of the Douge TMP. Pink helices correspond to the deposited model, while green helices represent unmodelled helical densities. **d.** 3D reconstruction showing the density corresponding to the Bxb1 TMP. The inset highlights the C-terminal end of a TMP subunit bending towards the capsid. **e.** Ribbon representation of the Bxb1 TMP. Pink helices correspond to the deposited model, while helices in other colors represent unmodelled helical densities.

**Supplementary Fig. 13.**
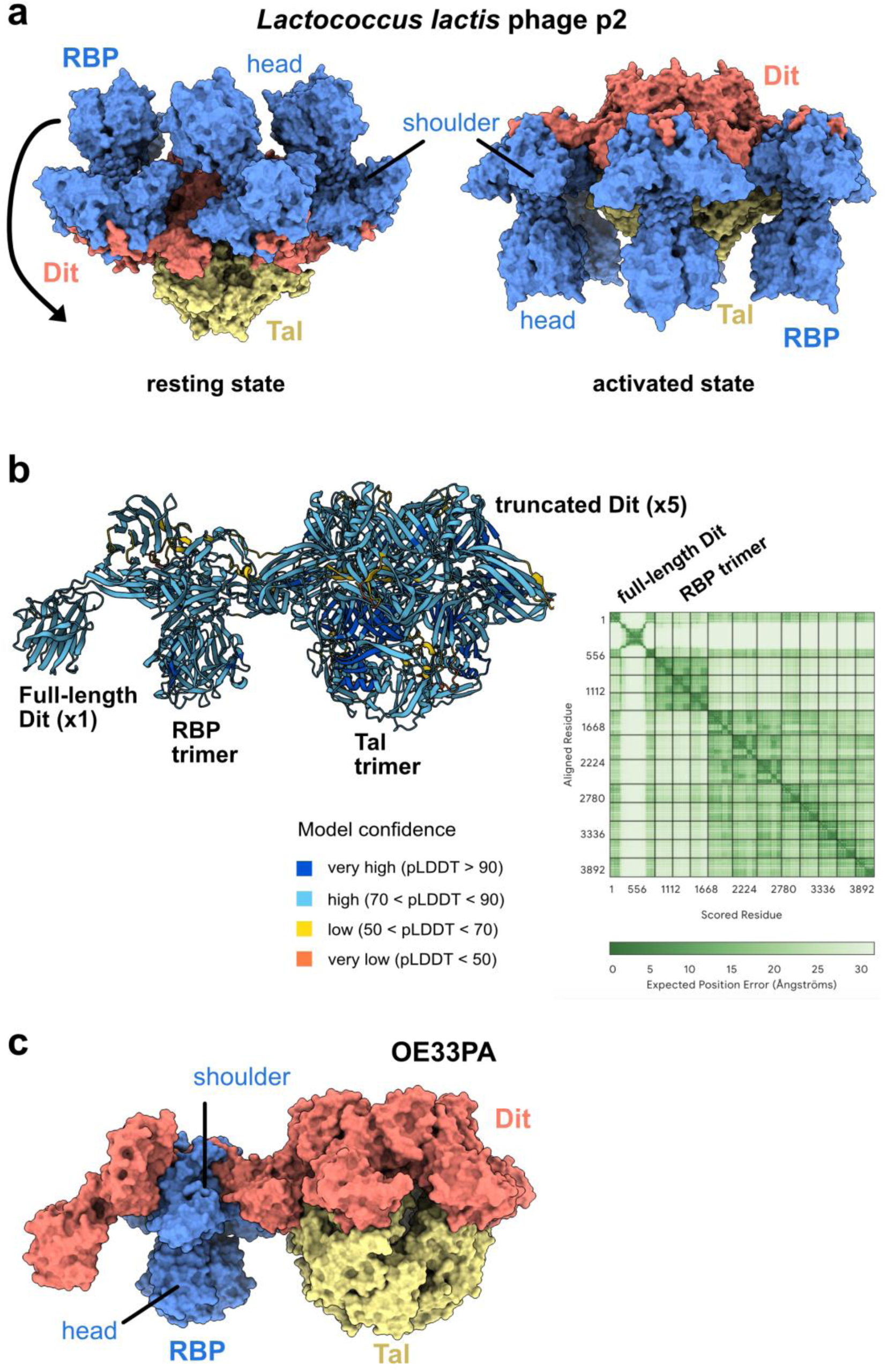
Conformational changes of the phages p2 and OE33PA adhesion device. **a.** Surface representation of the phage p2 adhesion device with the RBP head domains in an upward conformation (resting state, PDB ID 2wzp), and in a downward conformation (activated state, PDB ID 4v5i). The arrow suggests the rotation movement of the RBP upon activation. **b.** The AlphaFold3- predicted structure of a partial OE33PA adhesion device is shown as ribbons colored according to the pLDDT values. Because of calculation limitations, we have predicted the structure of only one RBP trimer and one full-length Dit. For the other 5 truncated Dit subunits, we replaced the three-domain extension (residues 147-537) by a GSGSG linker. The PAE plot shows confident interactions between the N-terminal end of the RBP trimer, the shoulder domain, and the Dit arm. **c.** Surface representation of the AlphaFold3-predicted structure shown in b.

**Supplementary Fig. 14.**
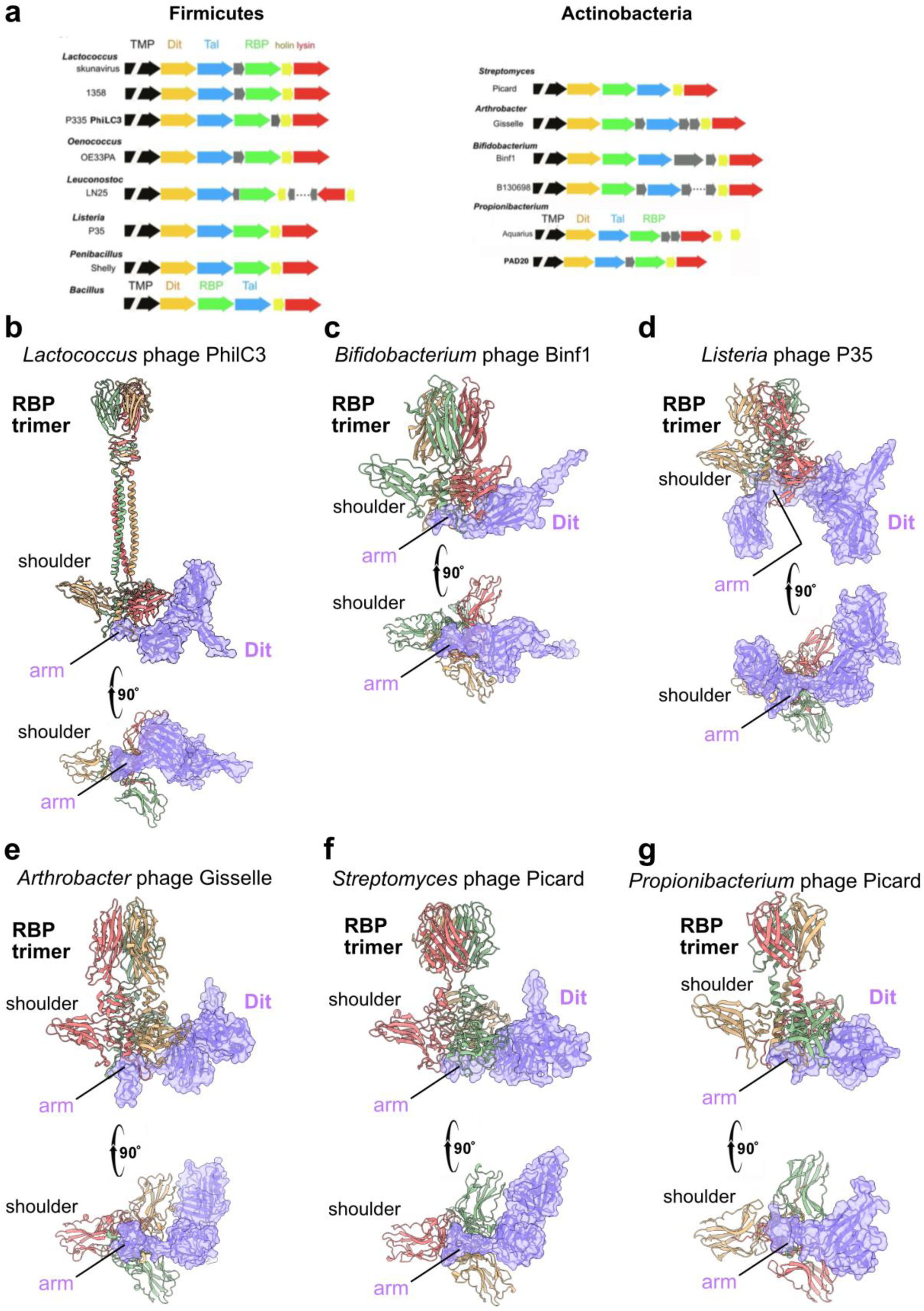
Candidate phages with an “articulated” adhesion device. **a.** Schematical representation of the ORF forming their adhesion device. All Tal are short. **b-g.** Orthogonal views of the AlphaFold3-predicted Dit–RBP complex for candidate “articulated” phages, showing RBP as a ribbon representation and Dit as both ribbon and surface representations. All Dit arm domains interact with the RBP shoulder domains.

**Supplementary Table 1.**
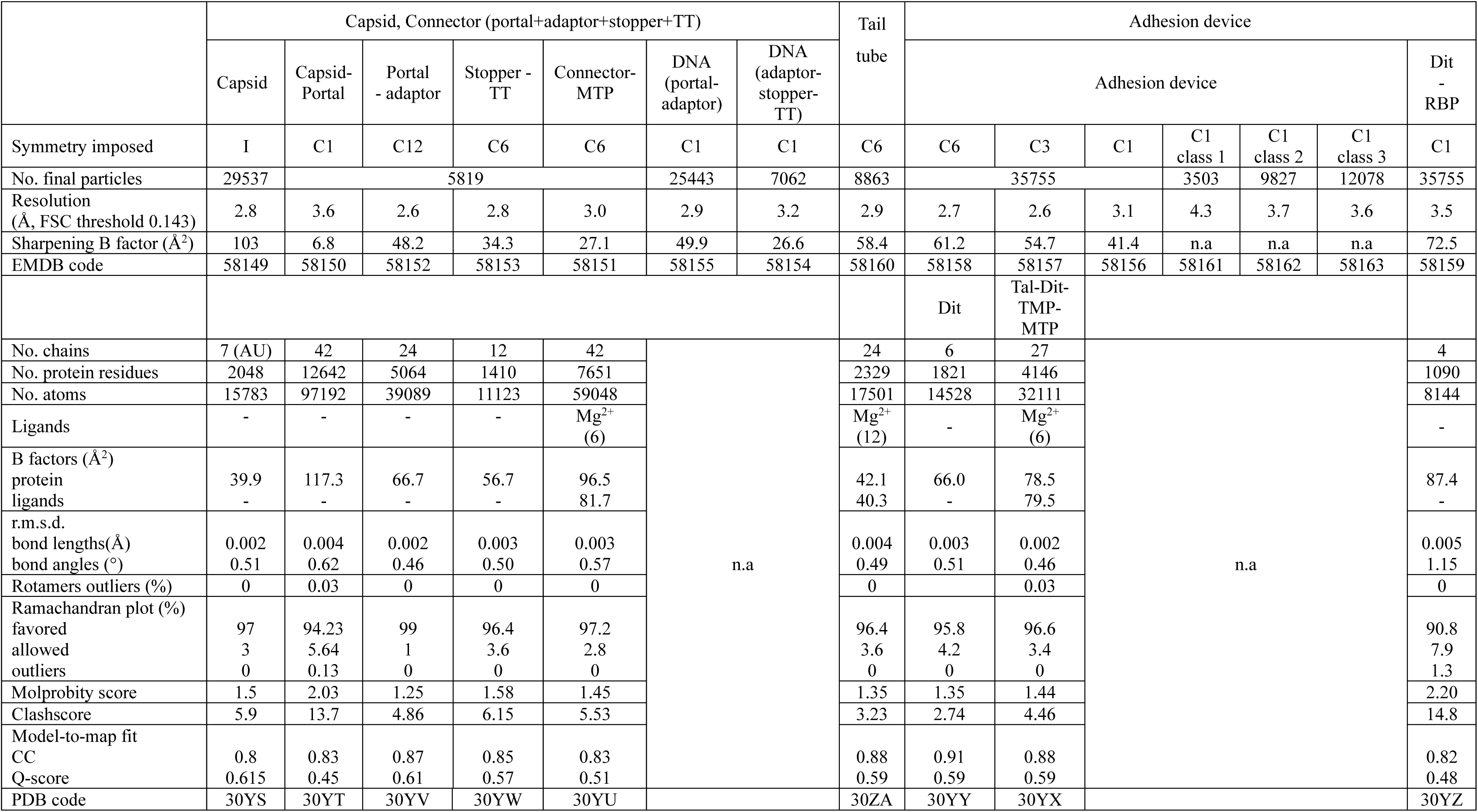
CryoEM 3D reconstruction and model refinement statistics.

|  | Capsid, Connector (portal+adaptor+stopper+TT) |  |  |  |  |  |  | Tail tube | Adhesion device |  |  |  |  |  |  |
|---|---|---|---|---|---|---|---|---|---|---|---|---|---|---|---|
|  | Capsid | Capsid-Portal | Portal - adaptor | Stopper - TT | Connector-MTP | DNA (portal-adaptor) | DNA (adaptor-stopper-TT) |  | Adhesion device |  |  |  |  |  | Dit - RBP |
| Symmetry imposed | I | C1 | C12 | C6 | C6 | C1 | C1 | C6 | C6 | C3 | C1 | C1 class 1 | C1 class 2 | C1 class 3 | C1 |
| No. final particles | 29537 | 5819 |  |  |  | 25443 | 7062 | 8863 | 35755 |  |  | 3503 | 9827 | 12078 | 35755 |
| Resolution<br>(Å, FSC threshold 0.143) | 2.8 | 3.6 | 2.6 | 2.8 | 3.0 | 2.9 | 3.2 | 2.9 | 2.7 | 2.6 | 3.1 | 4.3 | 3.7 | 3.6 | 3.5 |
| Sharpening B factor (Å<sup>2</sup>) | 103 | 6.8 | 48.2 | 34.3 | 27.1 | 49.9 | 26.6 | 58.4 | 61.2 | 54.7 | 41.4 | n.a | n.a | n.a | 72.5 |
| EMDB code | 58149 | 58150 | 58152 | 58153 | 58151 | 58155 | 58154 | 58160 | 58158 | 58157 | 58156 | 58161 | 58162 | 58163 | 58159 |
|  |  |  |  |  |  |  |  |  | Dit | Tal-Dit-TMP-MTP |  |  |  |  |  |
| No. chains | 7 (AU) | 42 | 24 | 12 | 42 | n.a |  | 24 | 6 | 27 | n.a |  |  |  | 4 |
| No. protein residues | 2048 | 12642 | 5064 | 1410 | 7651 |  |  | 2329 | 1821 | 4146 |  |  |  |  | 1090 |
| No. atoms | 15783 | 97192 | 39089 | 11123 | 59048 |  |  | 17501 | 14528 | 32111 |  |  |  |  | 8144 |
| Ligands | - | - | - | - | Mg<sup>2+</sup><br>(6) |  |  | Mg<sup>2+</sup><br>(12) | - | Mg<sup>2+</sup><br>(6) |  |  |  |  | - |
| B factors (Å<sup>2</sup>)<br>protein<br>ligands | 39.9<br>- | 117.3<br>- | 66.7<br>- | 56.7<br>- | 96.5<br>81.7 |  |  | 42.1<br>40.3 | 66.0<br>- | 78.5<br>79.5 |  |  |  |  | 87.4<br>- |
| r.m.s.d.<br>bond lengths(Å)<br>bond angles (°) | 0.002<br>0.51 | 0.004<br>0.62 | 0.002<br>0.46 | 0.003<br>0.50 | 0.003<br>0.57 |  |  | 0.004<br>0.49 | 0.003<br>0.51 | 0.002<br>0.46 |  |  |  |  | 0.005<br>1.15 |
| Rotamers outliers (%) | 0 | 0.03 | 0 | 0 | 0 |  |  | 0 |  | 0.03 |  |  |  |  | 0 |
| Ramachandran plot (%)<br>favored<br>allowed<br>outliers | 97<br>3<br>0 | 94.23<br>5.64<br>0.13 | 99<br>1<br>0 | 96.4<br>3.6<br>0 | 97.2<br>2.8<br>0 |  |  | 96.4<br>3.6<br>0 | 95.8<br>4.2<br>0 | 96.6<br>3.4<br>0 |  |  |  |  | 90.8<br>7.9<br>1.3 |
| Molprobity score | 1.5 | 2.03 | 1.25 | 1.58 | 1.45 |  |  | 1.35 | 1.35 | 1.44 |  |  |  |  | 2.20 |
| Clashscore | 5.9 | 13.7 | 4.86 | 6.15 | 5.53 |  |  | 3.23 | 2.74 | 4.46 |  |  |  |  | 14.8 |
| Model-to-map fit<br>CC<br>Q-score | 0.8<br>0.615 | 0.83<br>0.45 | 0.87<br>0.61 | 0.85<br>0.57 | 0.83<br>0.51 |  |  | 0.88<br>0.59 | 0.91<br>0.59 | 0.88<br>0.59 |  |  |  |  | 0.82<br>0.48 |
| PDB code | 30YS | 30YT | 30YV | 30YW | 30YU |  |  | 30ZA | 30YY | 30YX |  |  |  |  | 30YZ |

**Supplementary Table 2.** Similarity between the structures of the OE33PA virion components and PDB deposited structures.

| OE33PA<br>Protein | Hit § | E-value § | Seq.<br>id. % § | Rmsd<br>Å # | n. aligned<br>residues # | PDB<br>Id |
|---|---|---|---|---|---|---|
| Capsid | Native GTA particle | 2.1 e-17 | 18.7 | 3.3 | 255/297 | 6tb9 |
|  | Phage HK97 | 3.4 e-14 | 14.6 | 3.6 | 253/280 | 2fs3 |
| Portal | Native GTA article | 2.37 e-11 | 13.1 | 4.0 | 297/360 | 6te8 |
|  | Phage HK97 | 1.19 e-10 | 11.9 | 6.3 | 293/346 | 8fql |
| Adaptor | Phage HK97 | 1.02 e-2 | 11.7 | 2.0 | 90/96 | 3jvo |
|  | Phage SPP1 | 3.13 e-1 | 15.5 | 3.2 | 89/102 | 7z4w |
| Stopper | Native GTA particle | 8.31 e-4 | 20.9 | 3.6 | 101/110 | 6te9 |
|  | Phage SPP1 | 9.52 e-3 | 14.6 | 5.0 | 90/109 | 7z4w |
| Tail terminator | Myophage Milano | 4.94 e-5 | 9.0 | 3.2 | 110/155 | 8fwe |
|  | Phage SPP1 | 7.64 e-3 | 9.6 | 3.7 | 95/139 | 2lfp |
| MTP | Myophage E217 | 3.78 e-2 | 8.2 | 3.7 | 132/144 | 8fuv |
|  | Phage T5 | 1.51 e-1 | 14.4 | 3.6 | 137/440 | 5ngj |
| Dit | Phage p2 | 4.28 e-13 | 15.9 | 3.9 | 244/298 | 4v5i |
|  | Phage SPP1 | 7.35 e-10 | 10.2 | 3.0 | 227/242 | 2x8k |
| Tal | Phage p2 | 2.64 e-14 | 15.6 | 3.9 | 255/372 | 4v5i |
|  | Phage MC58 | 7.44 e-12 | 10.6 | 3.7 | 263/326 | 3d37 |
| RBP shoulder | Phage p2 | 1.21 e-9 | 17.6 | 3.8 | 143/254 | 2wzp |
|  | Phage 1358 | 6.03 e-8 | 16.3 | 3.0 | 216/390 | 4l92 |
| RBP head | Phage bIL170 | 1.14 e-4 | 23.2 | 2.1 | 99/110 | 2fsd |
|  | Phage TP901-1 | 7.53 e-4 | 19.3 | 2.8 | 92/99 | 4ios |

§ data from Foldseek ; # data from Dali server

